# Differential regulation of histone H1 subtypes by N6-methyladenosine RNA methylation

**DOI:** 10.1101/2025.01.22.634368

**Authors:** D García-Gomis, J López-Gómez, M Andrés, Z Fernández, A Benyahya, N Serna-Pujol, I Ponte, A Jordan, A Roque

**Author notes:** **Corresponding author**: Alicia Roque, Biochemistry and Molecular Biology Department, Biosciences Faculty, Universitat Autònoma de Barcelona, Spain.

## Abstract

Histone H1 is involved in the regulation of chromatin structure and gene expression. Up to seven H1 subtypes or variants are expressed in human somatic cells. The H1 complement, defined as the subtype composition and proportions in each cell, is highly variable depending on the cell type, cell cycle stage, developmental context, and several diseases such as cancer. This variability results from the combined action of multiple regulatory processes. Epitranscriptomic modifications have emerged as a new regulatory layer capable of controlling all aspects of mRNA metabolism. In this work, we examined the role of the most prevalent mRNA modification, N6 methyladenosine (m6A), in the regulation of H1 subtypes. MeRIP-seq showed that *H1-0* and *H1-4* are enriched in m6A, whereas *H1-2* displays intermediate levels of this mark. Integration of the functional studies involving m6A inhibition and partial depletion of these m6A readers led us to propose the first model of the differential regulation of H1 subtypes by m6A. In this model, m6A promotes the degradation of *H1-0* mRNA mediated by YTHDF2, while it enhances the stability of *H1-2* mRNA through IGF2BP1 binding and its translation by the combined action of IGF2BP1 and hnRNPD. In the case of *H1-4*, m6A promotes its transcription and stimulates its translation via hnRNPD binding. These findings suggest that m6A participates in the subtype-specific regulation of H1 variants, adding another layer to their complex regulation and contributing to the variability of the H1 complement in cancer.

**Keypoints:**

- Histone transcripts have different levels of m6A.
- We analyzed the function of m6A in *H1-0*, *H1-2*, and *H1-4* and identified its readers.
- We propose the first model of the role of m6A in the subtype-specific regulation of histone H1.

## Introduction

Histone H1 is involved in the regulation of chromatin structure and gene expression. In humans, Histone H1 is a protein family composed of eleven subtypes or variants. Somatic cells express up to seven subtypes, while the remaining four are germline-specific. H1 somatic subtypes are divided into two groups. The first group, the replication-dependent subtypes (RD), includes subtypes H1.1-H1.5, whose genes are in the histone cluster of chromosome 6. The second group, the replication-independent subtypes (RI), includes H1.0 and H1X (1).

The existence of multiple H1 subtypes raises the question of whether they are functionally redundant or differentiated. Experimental evidence accumulated in the field shows that while there is compensation among subtypes after knockout, their differential properties and continued selective pressure suggest some degree of functional specificity (2) The differential expression patterns determine the H1 complement, defined as the subtype composition and proportions in a specific cell. The H1 complement is variable in physiological conditions according to the cell type, cell cycle phase, and developmental stage (2). Alterations of the H1 complement have been described in disease. Changes in the mRNA and protein levels and some post-translational modifications have been associated with different cancer types (3–6). Therefore, studying the regulation of H1 subtypes is essential for understanding their specific functions and role in disease.

Coordinated action of several regulatory mechanisms at transcriptional, post-transcriptional, translational, and post-translational levels modulate the H1 complement. RD subtype transcription occurs mainly during the S-phase, as their genes are part of the histone locus body (HLB) (7). This subcellular membrane-less compartment is formed during S-phase and includes the histone cluster and all the transcriptional machinery. HLB also contains the components needed for the post-transcriptional processing of the RD transcripts, which results in the formation of a stem-loop at their 3’ end, without a polyA tail (8).

Histone H1 RI subtypes are encoded in chromosomes 3 and 22, which are not part of the HLB. They are transcribed throughout the cell cycle, although their mRNA levels are highest during S-phase. However, the *H1-0* transcript accumulates in cells arrested in G0 (9). *H1-0* and *H1-10 (H1X)* transcripts have longer untranslated regions (UTR) at both ends of the transcript than RD subtypes and their 3’UTR is polyadenylated (1). The transcript levels of all somatic subtypes are co-regulated, which explains at least in part the compensation observed after the partial or total depletion of one or more subtypes (10).

Transcript levels of H1 subtypes do not always correlate with the protein levels, suggesting that other molecular mechanisms are involved in the subtype-specific regulation (6, 10). We have recently shown that H1 subtypes have different stability at the protein level (11). However, other mechanisms might contribute to the control of their transcript levels and translation.

In recent years, it has been shown that base modifications in RNA, known as epitranscriptome modifications, can regulate RNA metabolism. The most abundant modification in the mRNA is the methylation of adenine at position 6, m6A (12). This modification is incorporated in the mRNA consensus sequence DRACH (D=A/G/U, R=A/G, H=A/C/U) by METTL3, using S-adenosyl methionine (SAM) as a donor of the methyl group. This RNA methyltransferase is the only m6A writer that modifies mRNA and the catalytic subunit of a complex formed by the heterodimer METTL3-METTL14 and the auxiliary proteins WTAP, CLL1, VIRMA, RBM15, and ZC3H13. There are two erasers of m6A, the demethylases ALKBH5 and FTO (13, 14).

The recognition of m6A is carried out mainly by three different protein families: YTHDFs, IGF2BPs, and hnRNPs. There are two forms of m6A recognition: direct binding to the modified residue and the “m6A switch” or indirect mechanism. In the latter case, m6A induces RNA unfolding, increasing the reader’s accessibility to single-stranded RNA (15). The proteins belonging to the YTHDF family are direct readers of m6A, while the members of the IGF2BP family can read m6A directly or indirectly, and hnRNPs are mostly indirect readers (13, 16–18). At the functional level, m6A can regulate several facets of RNA processing, including alternative splicing, export, stability, and translation (13, 14). The proteins of the m6A pathway are often dysregulated in solid and blood tumors (reviewed in (19–21)). The presence of m6A has been reported in several H1 transcripts (22). Therefore, it is reasonable to think that this mechanism may contribute to the regulation of H1 subtypes and explain some of its alterations in cancer.

In this work, we have studied the role of m6A in the regulation of somatic H1 subtypes using synchronized HeLa cells. The analysis of m6A RNA immunoprecipitation coupled with next-generation RNA sequencing (meRIP-seq) data showed that m6A was enriched in the transcripts of *H1-0* and *H1-4* and had medium levels in *H1-2*. We also studied the m6A readers bound to H1 transcripts and performed functional analysis in the presence of m6A inhibitors. Our results showed that in H1 transcripts m6A was recognized by different readers and had distinct functional effects. These findings suggest that the role of m6A in the regulation of histone H1 is subtype-specific and contributes to the variability of the H1 complement.

## Materials and methods

### Cell culture and treatments

All the cell lines were grown at 37°C and 5% CO_2_ in their specific culture media supplemented with 10% fetal calf serum (Gibco) and 1% penicillin-streptomycin (Ddbiolab). Human embryonic kidney 293T cells (HEK293T) and cervical carcinoma cells (HeLa) were cultured in DMEM Glutamax (Corning). After harvesting, cells were counted in an automated cell counter (Bio-Rad). For m6A inhibition, cells were treated with 25, 50, or 100 mM cycloleucine (TCI, A1063) for 24h or with 20 µM STM2457 (Merck, SML3360) for 48h.

### Cell synchronization

HeLa cells were synchronized by a double thymidine block (23). Plates were seeded at 25% confluency, and 2 mM thymidine was added. After 18h, the medium was removed, and the cells were washed with phosphate buffer saline (PBS). Fresh medium was added, and the cells were grown for 9h. Subsequently, 2 mM thymidine was added, and the cells were incubated for another 18h. Finally, the medium was removed, the cells were washed with PBS, and a complete medium was added. Sample collection was done as follows: 0 hours G1/S phase, 3 hours S phase, 9 hours G2/M phase, and 15 hours G1 phase. The proportion of cells in each phase of the cycle was determined using flow cytometry.

### Flow cytometry

To analyze cell cycle progression, one and a half million cells were fixed in 70% ethanol and incubated at -20 °C for at least 2h. Fixed cells were centrifuged at 200g for 5 min and resuspended in 3 ml of PBS. After 1 minute on ice, the cells were centrifuged at 200g for 5 min. Afterward, cells were resuspended in staining solution (0.1% Triton X-100, 100 µg/mL RNase A, 20 ng/mL propidium iodide in PBS) and incubated for 30 min in the dark. Stained cells were analyzed by flow cytometry using a FACSCalibur cytometer (BD bioscience) and CellQuest Pro software.

### Partial depletion by siRNA

HeLa cells were transfected with 25 nM of *IGF2BP1*, *hnRNPD*, and *YTHDF2* custom siRNAs (Eurofins) (Table S1) or a negative control siRNA (1022076, Qiagen) using METAFECTENE SI+ (Biontex). After 48 h, the cells were harvested, and the effectivity of the siRNA depletion was confirmed by RT-qPCR (Table S1). The effect of partial depletion on mRNA and protein levels was analyzed using RT-qPCR and Western blot, respectively.

### Immunoprecipitation of m6A-RNA (meRIP)

The protocol described by (24) was adapted to the peculiarities of H1 transcripts. Total RNA from 2.5·106 cells was extracted with RNAqueous™ Total RNA Isolation Kit (Invitrogen). Protein A Dynabeads (Thermo Fisher) were washed in IP buffer containing 10 mM Tris-HCl pH 7.4, 250 mM NaCl, and 0.1% NP-40. In each immunoprecipitation reaction, magnetic beads were mixed with 4 μg of antibody (anti-m6A or IgG, see Table S2) in the presence of 0.5 mg/mL bovine serum albumin (BSA) for 2h at 4 °C with orbital agitation. The beads were washed with IP buffer, and 150 µg/per sample of denatured total RNA and the RNAase inhibitor (RNAseOUT, Thermo Fisher) were added. An aliquot of each sample was stored and used as input. The mixture was incubated for 2h at 4 °C with shaking, followed by three washes with IP buffer for 5 min at 4 °C with shaking. Subsequently, two consecutive elution steps were carried out in IP buffer supplemented with 6.7 mM m6A (Biosynth, NM32281) for 1h at 4 °C. The eluted RNA was precipitated, dried, and resuspended in nuclease-free water. RNA was quantified using the fluorimeter Quantus (Promega). The immunoprecipitated RNA was used for RT-qPCR and RNA sequencing.

### RNA sequencing (RNAseq)

Total RNA (input) and m6A-immunoprecipitated RNA from HeLa cells synchronized in G1, S, and G2/M were analyzed by RNAseq, carried out at the Beijing Genomics Institute (BGI, China). Before sequencing, mRNA was enriched by rRNA depletion. RNA was fragmented and reversed transcribed using random hexamers. Second-strand synthesis was carried out, and the samples were subjected to end-repair and then 3’ adenylated. Adaptors were ligated to the ends of these 3’ adenylated fragments. Fragments were PCR-amplified, and PCR products were purified and selected with the Agencourt AMPure XP-Medium kit. The double-stranded PCR products were heat-denatured and circularized by the splint oligo sequence. The single-strand circle DNAs (ssCir DNA) were formatted as the final library and then quality-checked. The library was amplified to make DNA nanoballs. Pair-end sequencing was performed on the DNBSEQ-T7 platform, obtaining 100 bp reads. Biological triplicates from all samples were sequenced.

### RNAseq data analysis

FASTQ files containing the sequence of each strand were aligned to the human genome GRCh38.87 using Kallisto STAR run in the Biojupies cloud (25). The aligned reads, present in only one of the three replicates, and low abundance transcripts (less than one count per million reads) were filtered out. Analysis of differential expression of genes during the cell cycle was analyzed by comparing the RNA-seq results of the input samples from two consecutive cell cycle phases with the DEseq2 algorithm, using 1.5-fold change and an FDR-adjusted p-value below 0.05 as cut-off values (26). To estimate the content of m6A, the RNAseq data from the immunoprecipitated samples were compared with the input samples with the DEseq2 algorithm using the same cut-off values described above. After the analysis, transcripts in the immunoprecipitated samples were divided into three groups according to the fold change of the transcripts in the immunoprecipitated samples relative to the input: 1) high levels of m6A, transcripts increased above 1.5-fold with an FDR-adjusted p-value below 0.05; 2) moderate m6A levels, transcripts that increased or decreased below 1.5 fold; and 3) low m6A levels, transcripts decreased above 1.5 fold with an FDR-adjusted p-value below 0.05. Raw sequencing data and DEseq2 results are available in the Gene Expression Omnibus (GEO) database under the identifier GSE236863.

### RT-qPCR

Total RNA was purified with RNAqueous™ Total RNA Isolation Kit (Invitrogen), following the manufacturer’s instructions. Purified RNA was quantified, and 100 ng were retrotranscribed with iScriptTM cDNA synthesis kit (Bio-Rad) using random hexamers as primers. cDNA was amplified by qPCR using primers specific for each intended target (Table S1). The reaction mixture contained one ng of cDNA, 0.5 μM of each primer, and SYBR supermix (BioRad) in a final volume of 20 μL. The cycle parameters were as follows: one cycle of 95 °C for 3 min, 40 cycles of 95 °C for 10 s, and 60 °C for 30 s, followed by the melting curve (65 °C-95 °C). Negative controls without retrotranscriptase were included in every assay for each gene. Mean Cqs were calculated for all H1 subtypes and GAPDH, the housekeeping gene used as a control. Triplicates with a difference in Cq values > 0.5 cycles were rejected. Fold change was calculated using the ddCT method (27).

### Dot blot assay for m6A

Total RNA was purified with RNAqueous™ Total RNA Isolation Kit (Invitrogen) and quantified. The samples were denatured for 3 minutes at 95°C, loaded onto a positively charged nylon membrane (Roche), and UV-crosslinked in a Stratalinker® UV crosslinker 1800 (Agilent) in autocrosslink mode (120 mJ, 25 seconds) twice. The membrane was washed once in PBST (PBS 1x 0.02% Tween-20) for 5 minutes and blocked for 1 hour in 5% nonfat milk in PBST. The membrane was incubated overnight at 4°C with shaking with antibody against m6A (Table S2) diluted in PBST with 2.5% nonfat milk. It was washed three times with PBST for 5 minutes. The membrane was incubated for 1 hour in the secondary antibody (Table S2) diluted in PBST with 2.5% nonfat milk. The membrane was washed 3 times for 5 min with washing solution, developed with ECL Clarity (Bio-rad), and visualized on a Chemidoc imaging system (Bio-rad).

### Nuclear run-on

Nascent RNA was labeled with bromouridine (BrdU) and immunoprecipitated with a specific antibody, as described in (28). Four million cells per sample were lysed in 10 mM Tris-HCl pH 7.4, 10 mM NaCl, 3 mM MgCl2, 0.5% NP-40 for 5 min on ice and then centrifuged at 300g for 4 minutes at 4°C. The nuclei were washed in the same buffer and resuspended in 50 mM Tris-HCl pH 8.3, 0.1 mM EDTA, 5 mM MgCl2, and 40% glycerol. Transcription mixture containing 20 mM Tris-HCl pH 8.3, 5 mM MgCl2, 300 mM KCl, 4 mM DTT, 100 U RNAseOUT, 1 mM ATP, 1 mM GTP, 1 mM CTP, 0.5 mM BrUTP and 0.5 mM UTP for the labeled sample or 1 mM UTP for the negative control was added to the nuclei in a 3:2, v/v proportion. The reaction mixture was incubated at 30 °C for 1 hour in a rotating wheel. Transcription was stopped by immediately extracting RNA with the High pure RNA isolation kit (Roche). For immunoprecipitation, Protein A Dynabeads (Thermo Fisher) in IP buffer containing 10 mM Tris-HCl pH 7.4, 250 mM NaCl, and 0.1% NP-40 were incubated with 2µg of anti-BrdU monoclonal antibody (Table S2) and 0.5 mg/ml BSA for 2h at 4 °C with orbital shaking. The beads were washed and resuspended with IP buffer with 100 U RNAseOUT, and 7 μg of denatured RNA were added. The IP mixture was incubated for 2h at 4 °C with orbital shaking. Beads were washed 3 times with IP buffer for 5 minutes at 4 °C with shaking, and RNA extraction was performed with TRIzol (Thermo Fisher). Purified RNA was resuspended in nuclease-free water and analyzed by RT- qPCR. Nascent transcription was evaluated by the ΔΔCt method, comparing the immunoprecipitated RNA with the input in the labeled and control samples.

### Targeted profiling of ribosome occupancy

Ribosome occupancy of the translation start site was analyzed by RNA digestion coupled with RT-qPCR, as described in (29). One culture plate with about ten million HeLa cells per condition was used. Briefly, the culture media was eliminated by aspiration, and the cells were washed with cold PBS supplemented with 100 µg/mL cycloheximide (Sigma). The cells were separated from the plate by scraping in 1 mL of 20 mM Tris-HCl pH 7.4, 150 mM NaCl, 5 mM MgCl2, 1 mM DTT, 1 mg/ml cycloheximide, 1% Triton X-100, resuspended and frozen in liquid nitrogen. The samples were thawed on ice and the cells were lysed mechanically, passing the solution through a needle.

Cellular debris was removed by centrifugation at 16000xg for 15 min at 4°C. The supernatant was recovered and the RNA concentration measured with Quantus (Promega). Per each transcript and condition, 500 ng of RNA were digested with 1000 U of RNAse I (Fisher Scientific) and 10 U of TurboDNase (Invitrogen) in 10 mM Tris-HCl pH 7.6, 2.5 mM MgCl2, 0.5 mM CaCl2, for 1h at 4°C. Then, 750 µL of TRIzol were added to purify undigested RNA. The RNA protected by the ribosome binding was retrotranscribed using specific primers that bind near the translation start site of the analyzed transcripts and quantified by qPCR (Table S1).

### RNA immunoprecipitation (RIP)

Five million HeLa cells treated with m6A inhibitors or untreated were harvested, pelleted by centrifugation for 5 minutes at 400g, and washed once with cold PBS. The cell pellets were lysed with 0.5 mL of lysis buffer (150 mM NaCl, 50 mM TRIS pH 8, 2 mM EDTA, 0.5% NP-40, 0.5 mM DTT, 1:25 PIC, 1 U/μL RNAseOUT) on ice for 30 minutes. The lysates were centrifuged at 16000g for 20 minutes. An aliquot of 50 μL cell lysate was saved as input and mixed with 1 mL TRIzol for RNA extraction. Dynabeads™ Protein A (Thermo Fisher) were incubated with 4 µg of YTHDF2 antibody or control IgG (Table S2) for 1h at 4°C in TENT buffer in the presence of BSA 0.5 mg/mL. The beads were washed and resuspended in TENT buffer. At this point, 200 µL of the lysate were added and incubated by rotating for 2 hours at 4°C. After separation using a magnetic rack, beads were washed with ice-cold TENT buffer three times. Beads were mixed with 1 mL TRIzol and recovered for RNA or resuspended in protein loading buffer for Western blot. The immunoprecipitation of specific mRNAs was analyzed by RT-qPCR.

### Biotin pull-down assay

Transcripts corresponding to *H1-2* and *H1-4* were selected by pull-down using specific biotinylated probes (30). Streptavidin C1 Dynabeads ™ MyOne ™(Thermo Fisher) were incubated with 100 pmol of the specific or control biotinylated probes (Table S1) in TENT buffer (10 mM Tris-HCl pH 8, 1 mM EDTA, 150 mM NaCl, and 0.5% Triton X-100) supplemented with of 0.5 mg/ml BSA for 1h at 4°C. For each reaction, HeLa cells were lysed in 5 mM Tris HCl pH 7.6, 150 mM NaCl, 0.5% NP-40, 0.5% sodium deoxycholate, 0.1% SDS, cOmplete protease inhibitor cocktail (PIC, Roche), RNAseOUT for 30 min at 4°C. The extract was clarified by centrifugation at 16000xg for 20 min at 4°C. The supernatant was used as input for the pull-down. The beads were washed once with TENT buffer and 40 µg of the cell extract were added in TENT buffer with PIC and RNAseOUT and incubated for 1h at 4 °C with orbital shaking. Subsequently, three washes were performed in TENT buffer with orbital shaking. The proteins associated with the specific mRNAs were analyzed using mass spectrometry and western blot. In the case of proteomics samples, 2 washes were performed with 100 mM ammonium bicarbonate, and the proteins were digested on-bead with 10 ng/mL trypsin in 100 mM ammonium bicarbonate at 37 °C overnight. In the morning, fresh trypsin was added and left for another 4h at 37 °C. The supernatant containing the digested proteins was transferred to a new tube and formic acid was added to a final concentration of 5%. The samples were stored at -80°C.

### Mass spectrometry

The analysis was carried out at the Proteomics Facility UPF/CRG, Barcelona. Samples were desalted using a MicroSpin C18 column (Nest Group) prior to liquid chromatography mass spectrometry (LC- MS/MS) analysis. Samples were analyzed by using a mass spectrometer (Thermo Fisher) coupled to an EASY- nLC 1200 (Thermo Fisher). Peptides were loaded directly onto an analytical column and phase separated by reversed phase chromatography using a 50 cm column with a diameter of 75 μm, packed with 2 μm C18 particles. Chromatography gradients started with 95% solution A (0.1% formic acid in water) and 5% solution B (0.1% formic acid in 80% acetonitrile) at a flow rate of 300 nl /minute. It was gradually increased to 25% solution B (and 75% A) for 52 minutes and subsequently to 40% B in 8 minutes. After each analysis the column was washed for 10 minutes with solution B. The mass spectrometer was operated in positive ionization with a nanospray voltage of 2.4 kV and a temperature of 305 °C. The acquisition was done in data-dependent mode (DDA) and scanned with 1 microscan at resolution of 120,000 in a mass range of m/z 350-1400 with detection in the Orbitrap. Auto gain control (AGC) was set to standard and injection to auto. In each DDA cycle, followed by each scan, the most intense ions above the threshold (ion count at 10000) were selected for fragmentation. The number of precursor ions selected for fragmentation was determined by top speed acquisition algorithm and a one-minute dynamic exclusion. The spectra of the fragment ions were produced via high-energy collision dissociation (HCD) with a collision energy normalized to 28% and were acquired in the ion trap of the mass analyzer. The AGC and injection were set to standard and dynamic respectively and the isolation window to 1.4 m/z. BSA (New England biolabs) was analyzed between each sample to avoid crossover and ensure instrument stability, and the QCloud278 cloud was used to monitor longitudinal performance during the project. Proteome software. Discoverer suite (v2.5, Thermo Fisher Scientific) and the Mascot search engine search engine (v2.6, Matrix Science). The data were matched against the SwissProt human database (January 2022, 20,536 entries) and a list of common contaminants corresponding to decoy entries. An ion precursor mass tolerance of 7 parts per million for an MS1 level was used for identification. Trypsin was chosen as the enzyme and up to 3 miscleavages were allowed. The tolerance on the fragment ion mass was set to 0.5 Da for an MS2 spectrum. Methionine oxidation and N-terminal acetylation were used as variable modifications while cysteine carbamidomethylation was set as fixed modification. The false discovery rate (FDR) in peptide identification was set at a maximum of 5%. The SAINTexpress algorithm was used to score protein-protein interactions. comparing the proteins detected in the immunoprecipitated sample with the control and those in the sample with the specific probe (31). Proteins with an adjusted Bonferroni FDR (BFDR) less than 0.05 were considered positive hits. Triplicates of each sample were analyzed. The mass spectrometry proteomics data have been deposited to the ProteomeXchange Consortium via the PRIDE (32) partner repository with the dataset identifier PXD043530.

### Preparation of protein extracts

Total protein extracts were prepared as previously described (11). Cells were harvested by trypsinization and washed twice with PBS. For preparing total protein extracts, cells were resuspended in a 50 mM Tris pH 8 buffer containing 300 mM NaCl, 1 % Triton X-100, 0.5 % sodium deoxycholate, 3mM DTT, and a protease inhibitor cocktail (PIC), and incubated for 30 minutes on ice. The mixture was centrifugated at 16000xg, 30 minutes, at 4 °C. The suspension was passed through a needle until homogeneity and centrifugated at 16000xg, 30 minutes, at 4 °C. The supernatant contained a mixture of cellular proteins. All the protein extracts were analyzed by SDS-PAGE and stored at -20°C. Protein concentration was determined with Bradford.

### Western blot

Equivalent amounts of total proteins (20 μg) were separated in a 12% denaturing polyacrylamide gel electrophoresis (SDS-PAGE) and electrotransferred to a polyvinylidene difluoride membrane (PVDF) (EMD Millipore) at 100V for 1h. Immunoblot analyses were performed with the conditions recommended by the manufacturer for the primary and secondary antibodies (Table S2). The specificity of primary H1 antibodies has been validated using knock-down cell lines (33). Blots were visualized with Clarity Western ECL substrate (Bio-Rad) in a Chemidoc imaging system (Bio-Rad). Band intensities were quantified using Image Lab software (Bio-Rad). α-tubulin, GAPDH, or histone H3 were used as loading controls.

## Results

### Analysis of m6A levels during cell cycle

The epitranscriptome mark m6A is involved in all the steps of RNA metabolism. We wanted to know the relative levels of this mark in H1 transcripts and its role in their regulation. Histone H1 subtypes mRNA levels are variable during the cell cycle. Therefore, we examined m6A levels in H1 transcripts and their fluctuations in the cell cycle. We selected HeLa cells as a model, as this cell line is easily synchronized using a double thymidine block. We obtained three cell populations with approximately 80% enrichment in G1, S, and G2M, respectively (Figure S1). With these samples, we performed RNA-seq and m6A immunoprecipitation of the transcripts, followed by RNA-seq. First, we characterized the input samples by RNAseq and analyzed the changes in expression during the cell cycle. We found that from G1 to S-phase, 793 genes were upregulated, while 692 and 195 genes were upregulated from S to G2/M and from G2/M to G1, respectively (Figure S2A; Table S3). In all cases, the upregulated genes were enriched in GO terms associated with the molecular processes typical of the appropriate phase, confirming the homogeneity of the synchronized samples (Figure S2B).

Transcripts with m6A in each RNA sample from synchronized cells were selected by immunoprecipitation with a specific antibody against this modification and analyzed by RNAseq. The enrichment in m6A of each transcript was examined by comparing its abundance in the input samples and after immunoprecipitation. We could compare both samples because the immunoprecipitation was performed using unfragmented RNA. The differential expression analysis between the input and the immunoprecipitated sample of each phase of the cell cycle allowed us to determine the relative levels of m6A in each transcript. The transcripts enriched in the immunoprecipitated samples corresponded to those with high levels of m6A, those unchanged corresponded to transcripts with medium levels of m6A, and the ones showing lower abundance in the immunoprecipitated sample corresponded to those with low levels of m6A. However, using this experimental design were unable to determine the localization of m6A within the transcript.

Analysis of RNAseq data of the immunoprecipitated samples showed that m6A was present in more than 17000 transcripts, including protein-coding and noncoding RNAs. We found that 5713 transcripts had high levels of m6A in G1, while 5979 and 5137 transcripts were enriched in this mark in S-phase and G2/M, respectively (Figures 1A and 1B; Table S4). Interestingly, almost 70% of the genes with high levels of m6A were conserved in all phases, as shown in the Venn diagram, suggesting that this modification remains stable during the cell cycle (Figure 1C). The GO term analysis showed that the transcripts with high m6A levels during the cell cycle encode proteins involved in key biological processes such as regulation of macromolecule biosynthesis, transcription, RNA processing, cell-to-cell adhesion, and regulation of gene expression (Figure 1D).

**Figure 1.**
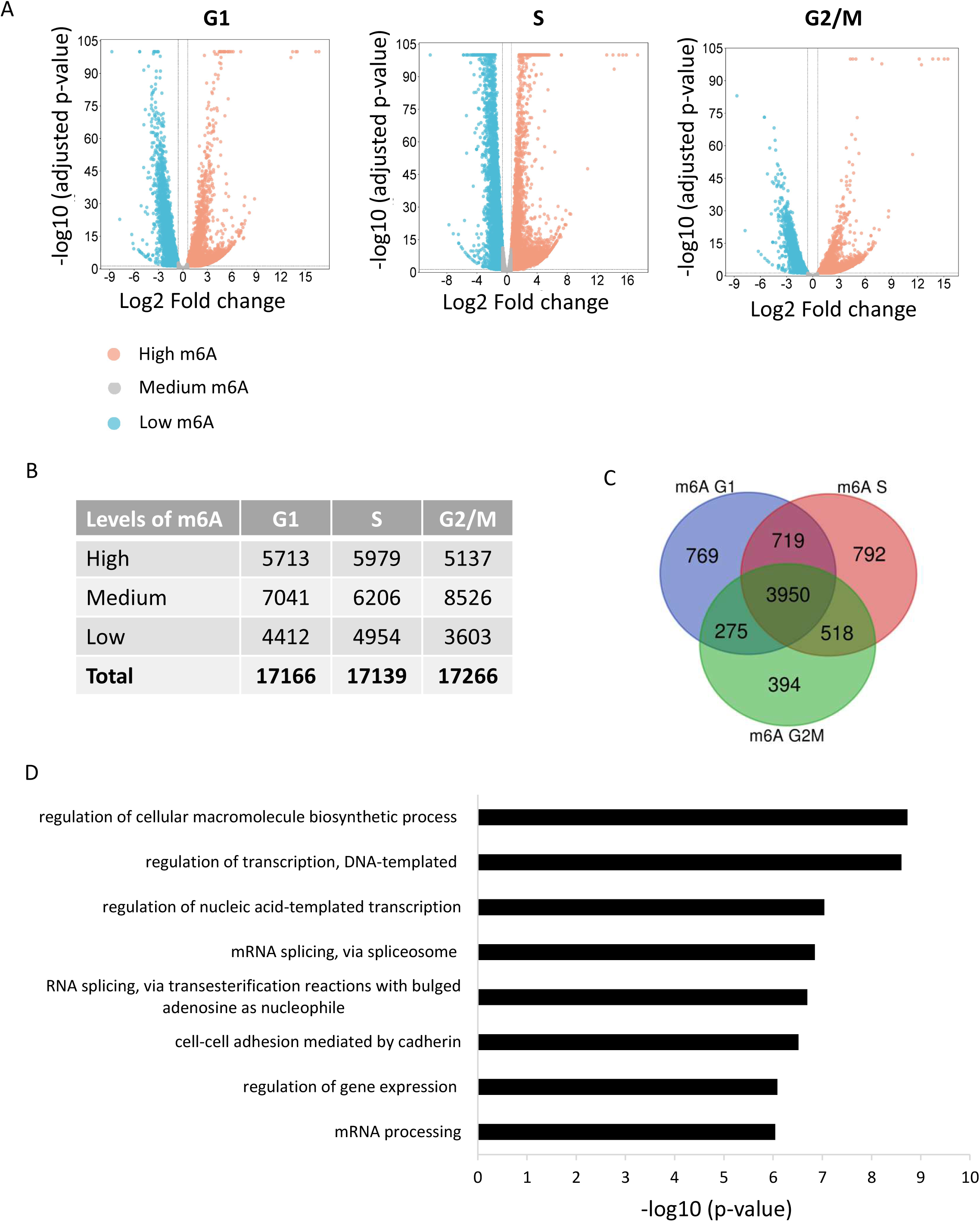
m6A levels during HeLa cell cycle. A. Vulcano plots of the DESeq2 analysis comparing the transcript levels in the m6A immunoprecipitated samples in each phase of the cell cycle relative to the input. The transcripts were divided into three groups: 1) high levels of m6A, transcripts increased above 1.5-fold in the IP samples with FDR-adjusted p-value <0.05; 2) medium m6A levels, transcripts that increased or decreased below 1.5-fold in the IP samples; and 3) low m6A levels, transcripts decreased above 1.5-fold in the IP samples with FDR-adjusted p-value <0.05. B. Results of the DESeq2 analysis. C. Venn diagram of the transcripts with high levels of m6A in each phase of the cell cycle. D. GO term analysis showing the biological processes enriched in the genes whose transcripts have high levels of m6A in all cell cycle phases.

### Expression and m6A levels of H1 subtypes during cell cycle

As our main objective was to study if m6A plays a role in the regulation of histone H1 subtypes, we analyzed their expression and m6A levels during the cell cycle. The RNAseq analysis of the input samples showed that six H1 subtypes are expressed in HeLa cells (Figure 2A). Subtype *H1-5* had the highest transcript levels, closely followed by *H1-2* and *H1-10* (H1X). Subtype *H1-4* had medium mRNA levels, while *H1-0* and *H1-1* had the lowest expression. H1 subtypes were also differentially expressed during the cell cycle (Figure 2B). Three different patterns were observed. Replication-dependent subtypes were upregulated in S-phase and downregulated in G2/M. Subtype *H1-0* was upregulated in the S-phase and downregulated in G2/M and G1, while *H1-10* (H1X) was slightly upregulated in the S-phase, remained unchanged during G2/M, and was downregulated in G1.

**Figure 2.**
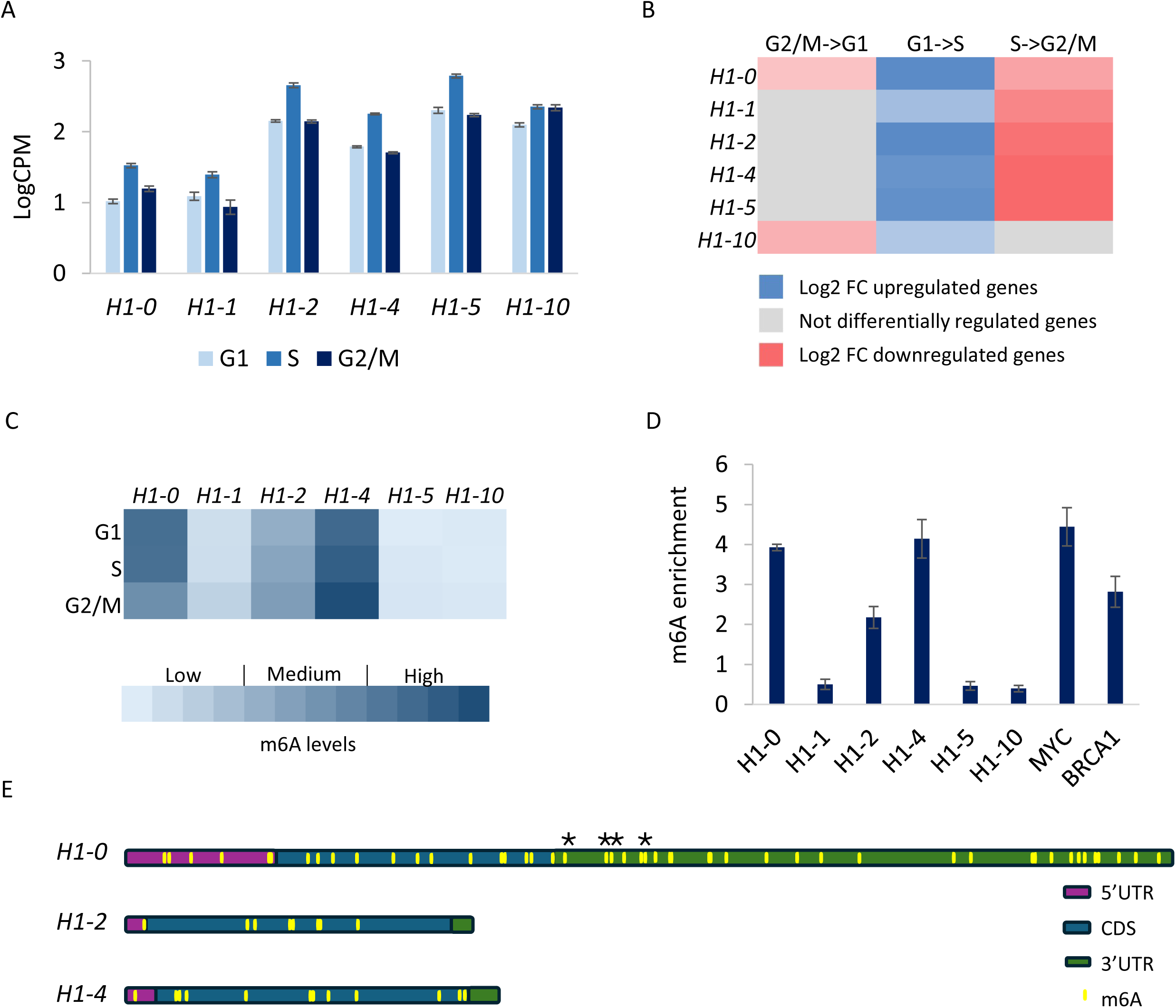
Expression changes and m6A levels of H1 subtypes during HeLa cell cycle. A. Normalized expression of H1 subtypes obtained by RNAseq of synchronized HeLa cells. B. Differential expression between cell cycle phases of H1 subtypes analyzed by DESeq2. Shades of red and blue are proportional to the log2FC values of the differentially expressed genes. C. m6A levels of H1 subtypes obtained by meRIPseq analysis. D. m6A enrichment in HeLa asynchronous cells analyzed by m6A-RIP-RT-qPCR. m6A enrichment values correspond to the fold change of the gene-specific RT-qPCR of the immunoprecipitated sample with respect to the input. *MYC* and *BRCA1* were used as positive control genes. E. m6A sites in human *H1-0*, *H1-2*, and *H1-4* transcripts are listed in RMBase v3.0 (22). Transcript regions and mRNA length are drawn to scale. Error bars correspond to the standard deviation of three biological replicates. CDS, coding sequence, CPM, counts per million. FC, fold change. RIP, RNA immunoprecipitation. RT, retrotranscription. qPCR, quantitative PCR. UTR, untranslated region.

The m6A levels of H1 transcripts were heterogeneous (Figure 2C). According to our meRIPseq data, the *H1-0* transcript had high levels of m6A in G1 and S, but they decreased to medium levels during G2/M. The levels of m6A in the other H1 transcripts remained in the same category during the cell cycle, with *H1-4* having high levels of m6A, *H1-2* medium levels, and low levels for the rest of the subtypes. The analysis by RT-qPCR of H1 subtypes in asynchronous HeLa RNA samples before and after m6A-specific-IP confirmed the levels of m6A enrichment obtained in the meRIPseq analysis (Figure 2D). The levels of m6A present in *H1-0* and *H1-4* were comparable to those obtained for *MYC* and *BRCA1* transcripts, which were detected as transcripts with high m6A levels by meRIPseq and are regulated by this mark (20, 34–37). Considering these results, we focused on the H1 subtypes with medium or high levels of m6A and performed the rest of the experiments using asynchronous cells.

As we previously explained, using our modified meRIP protocol, we cannot determine the localization of m6A within H1 transcripts. We explored previous analysis of m6A performed in human cells publicly available at RMBase v3.0 (22). In this database, there are 48 m6A sites reported for *H1-0*, 12 for *H1-4*, and 8 for *H1-2* (Figure 2E, Table S5). In the *H1-0* transcript, about 60% of the m6A sites are in the 3’UTR, while in *H1-2* and *H1-4*, they are mostly in the coding region. Nucleotide-resolution mapping of m6A sites in poly-A enriched HeLa transcripts using eTAM-seq (evolved TadA-assisted N6-methyladenosine sequencing) detected four positions with m6A in the H1-0 mRNA in the 3’UTR, close to the stop codon, confirming the presence of the modification (Figure 2E) (38).

### Role of m6A in the regulation of H1 subtypes

To study the effects of m6A on H1 subtypes, we used two different inhibitors: cycloleucine, which prevents the biosynthesis of S-adenosylmethionine (SAM)—the primary methyl donor for m6A deposition, and STM2457, a specific inhibitor of METTL3, the only known enzyme capable of incorporating m6A (20, 39–43). The treatment conditions for both inhibitors had a slight effect on cell growth, while the reduction of the total m6A levels was confirmed by dot blot (Figure S3).

The presence of m6A can regulate different aspects of mRNA metabolism, including its stability, degradation, and translation. Thus, we explored how its inhibition affected the levels of mRNA, de novo transcription, ribosome occupancy, and protein levels. Treatment with STM reduced the levels of m6A in the transcript of *H1-0*, *H1-2*, and *H1-4* to about 30%, 70%, and 50% of the amount in the untreated cells. A similar effect was observed in *BRCA1*, which is known to be regulated by m6A (34) (Figure 3A). The analysis of the mRNA levels by RT-qPCR after treatment with STM showed an over two-fold increase of *H1-0,* a 60% decrease of *H1-2,* and similar values for *H1-4* (Figure 3B). As expected, *BRCA1* mRNA levels also increased upon m6A inhibition (34). These results suggest that m6A could regulate the transcript levels of H1 subtypes by promoting the degradation of *H1-0* and the stability of *H1-2*.

**Figure 3.**
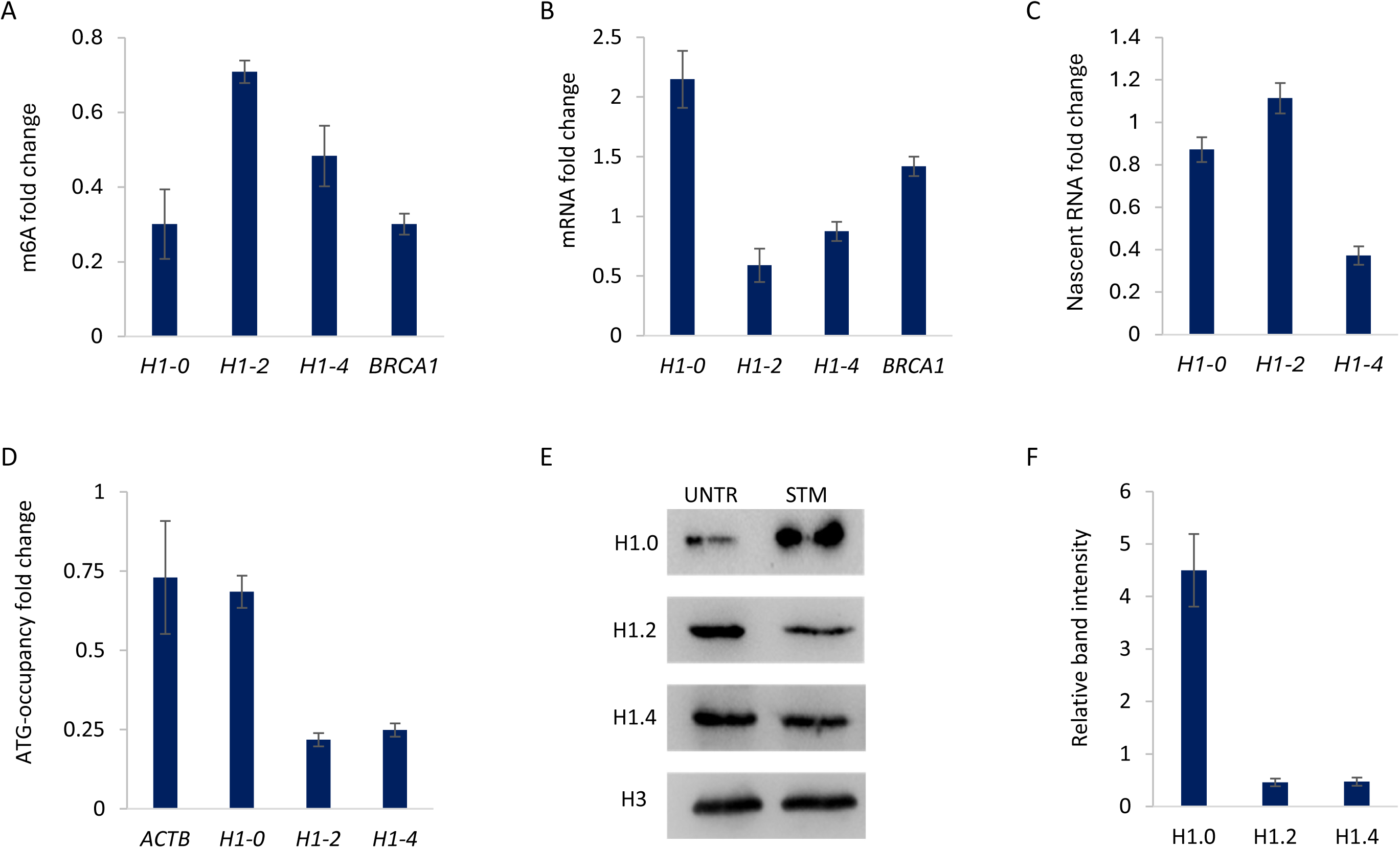
Effects of m6A inhibition by treatment with STM2457 (STM). A. Fold change of the m6A-immunoprecipitated mRNA. B. Fold change of the mRNA levels. C. Fold change of the BrdU-labeled nascent RNA immunoprecipitated after nuclear run on. D, Fold change of the transcript levels of the translation start-site region obtained by targeted ribosome profiling. E, Western blots of total protein extracts. F, Quantification of Western blot images. HeLa cells were treated with 20 μM STM for 48h. Fold change was calculated respect to the untreated cells (UNTR), which were used as a negative control in all the experiments. Error bars correspond to the standard deviation of three biological replicates.

The alterations in the mRNA levels could arise from changes in nascent transcription and in mRNA stability. Therefore, we analyzed nascent transcription by nuclear run-on in the presence of bromouridine. Our results showed no changes in the levels of nascent transcripts upon m6A inhibition for *H1-0* and *H1-2*, with values of fold change close to one, while those of *H1-4* decreased to less than a 40% of the untreated cells (Figure 3C). These results, combined with the changes in the mRNA levels, suggest that m6A has a differential role in the regulation of the three H1 subtypes.

The levels of m6A regulate the protein levels directly by promoting translation. We analyzed changes in the ribosome occupancy of the mRNA region surrounding the translation start site (ATG) of *H1-0*, *H1-2*, and *H1-4* by specific RT-qPCR (29). The mRNA levels of the first ribosome footprint after m6A inhibition were around 80% those in the untreated cells for *H1-0* and *ACTB* (β-actin), which was used as a control (Figure 3D). In contrast, the mRNA levels surrounding the initiation codon of *H1-2* and *H1-4* markedly decreased upon m6A inhibition, suggesting that m6A plays a role in the translation of both subtypes.

Next, we analyzed whether the changes in the mRNA levels and the occupancy of the translation start site affected the protein levels of the H1 subtypes by Western blot (Figures 3E and 3F). After m6A inhibition, the protein levels of H1.0 increased about four-fold, while H1.2 and H1.4 decreased more than 50%. These results showed that the changes in the protein levels of the three H1 subtypes reflect the m6A-dependent alterations observed in mRNA metabolism.

Treatment with cycloleucine caused changes in the mRNA and protein levels of H1 subtypes with the same tendency as that observed with STM, although the magnitude was different (Figure S4 and 3). We also observed the reduction of transcription and translation of H1-4 mRNA but failed to detect the reduction in the translation of H1-2 (Figure S4C, S4D, 3C, and 3D). The differences between the effects of the treatment with STM and cycloleucine probably arise from indirect effects of the latter.

We also inhibited m6A with cycloleucine in HEK293T (Figure S5). This cell line was more sensitive to this inhibitor, so we reduced its dose to 25 mM, which did not compromise cell viability and reduced m6A (Figures S5A and S5B). The analysis by RT-qPCR showed that the effect of cycloleucine on the mRNA levels was like the one observed with STM in HeLa (Figures S5C and 3B). The protein levels examined by Western blot also confirmed the results obtained in HeLa (Figures S5D, S5E, 3E, and 3F). Interestingly, the increase of H1.0 was higher in HEK293T than in HeLa (Figures S5D, S5E, 3E, and 3F). Overall, the changes described in HeLa were also observed in HEK293T, suggesting that the role of m6A in the regulation of histone H1 subtypes is conserved, at least in both cell types.

### Identification of m6A readers involved in the regulation of H1 subtypes

The functional role of m6A is associated with its recognition by different readers. We performed a specific mRNA pull-down to identify which m6A readers were bound to the transcripts of *H1-2* and *H1-4* by mass spectrometry. Specific biotinylated probes for both transcripts and a control probe, which was not complementary to any human transcript, were designed (Table S1). After the pull-down, we analyzed, by RT-qPCR, the transcripts of the RD H1 subtypes expressed in HeLa and *GAPDH* and *ACTB(*β-actin) (control genes) (Figure S6). We observed that each probe enriched the preparation in the desired transcript by more than 3.5-fold compared with the other H1 subtypes and the two control genes, indicating probe specificity. The proteins bound to each probe were identified by mass spectrometry, and the proteins differentially enriched in the pull-downs were selected for further analysis.

We found 36 and 31 proteins significantly enriched in the *H1-2* and *H1-4* transcript pull-down, respectively (Tables S6 and S7). All the identified proteins but one were mRNA-binding proteins, indicating that the pull-down was successful. Functional analysis using the STRING database showed that, in both cases, the identified proteins formed a network significantly enriched in protein-protein interactions (PPI) (Figures 4A and 4B). Gene ontology enrichment analysis revealed similar biological processes enriched in both interaction networks, including the mRNA metabolic process, stress granule assembly, and post-transcriptional regulation of gene expression (Figure 4C and 4D). This similarity is determined by the 17 proteins common to both networks, which are involved in the biological processes mentioned above (Figures S7A and S7B). We also confirmed the proteomic results by Western blot, detecting the presence of one shared protein, hnRNPM, in the pull-downs, while it was absent in the control IgG (Figure S7C).

**Figure 4.**
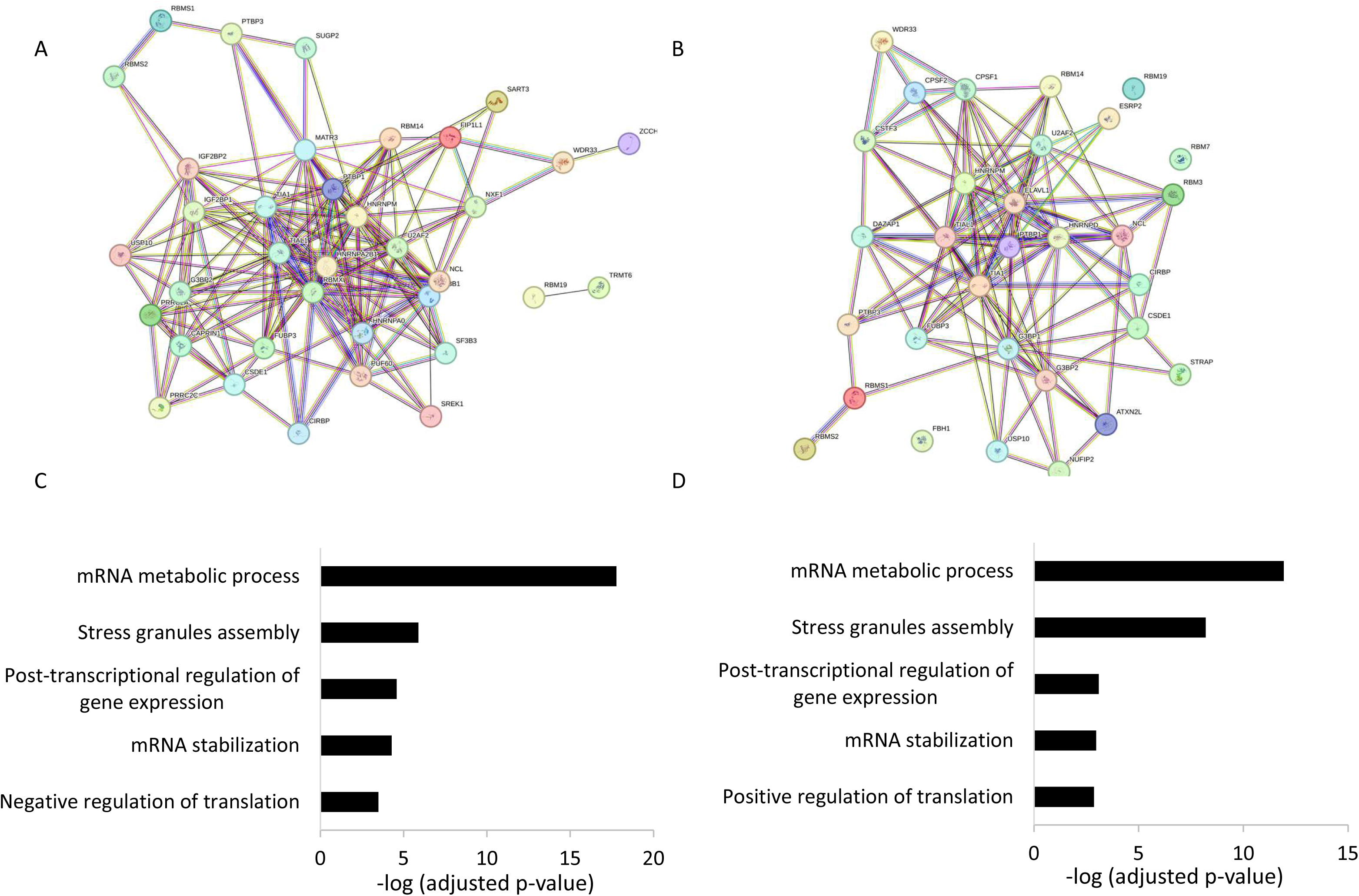
Proteomic analysis of the proteins bound to H1 transcripts. A, C. Interaction network obtained using STRING database for the proteins bound to *H1-2* and *H1-4* transcripts, respectively. Both networks have a PPI enrichment p-value: < 1.0e-16. B, D. Significantly enriched biological processes in the interaction networks formed by the proteins bound to *H1-2* and *H1-4* transcripts, respectively.

Specific proteins bound to the *H1-2* transcript, although present in all GO terms, are determinants in mRNA stabilization and negative regulation of translation (Figure 4C). Two m6A readers were in those GO terms, IGF2BP1 and IGFBP2. Both readers are associated with mRNA stabilization, which seems to be the functional role of m6A in the regulation of *H1-2*. However, considering that IGF2BP1 was more enriched than IGF2BP2, we hypothesized that this was the main m6A reader responsible for the *H1-2* transcript stabilization. Western blot analyses showed that IGF2BP1 was enriched in the *H1-2* pull-down and undetectable in that of *H1-4*, confirming the proteomic results (Figure 5A). In the presence of STM and cycloleucine, we observed a decrease in the amount of IGF2PB1 in the pull-down, showing that its interaction with *H1-2* is mediated by m6A (Figure 5A and S8A).

**Figure 5.**
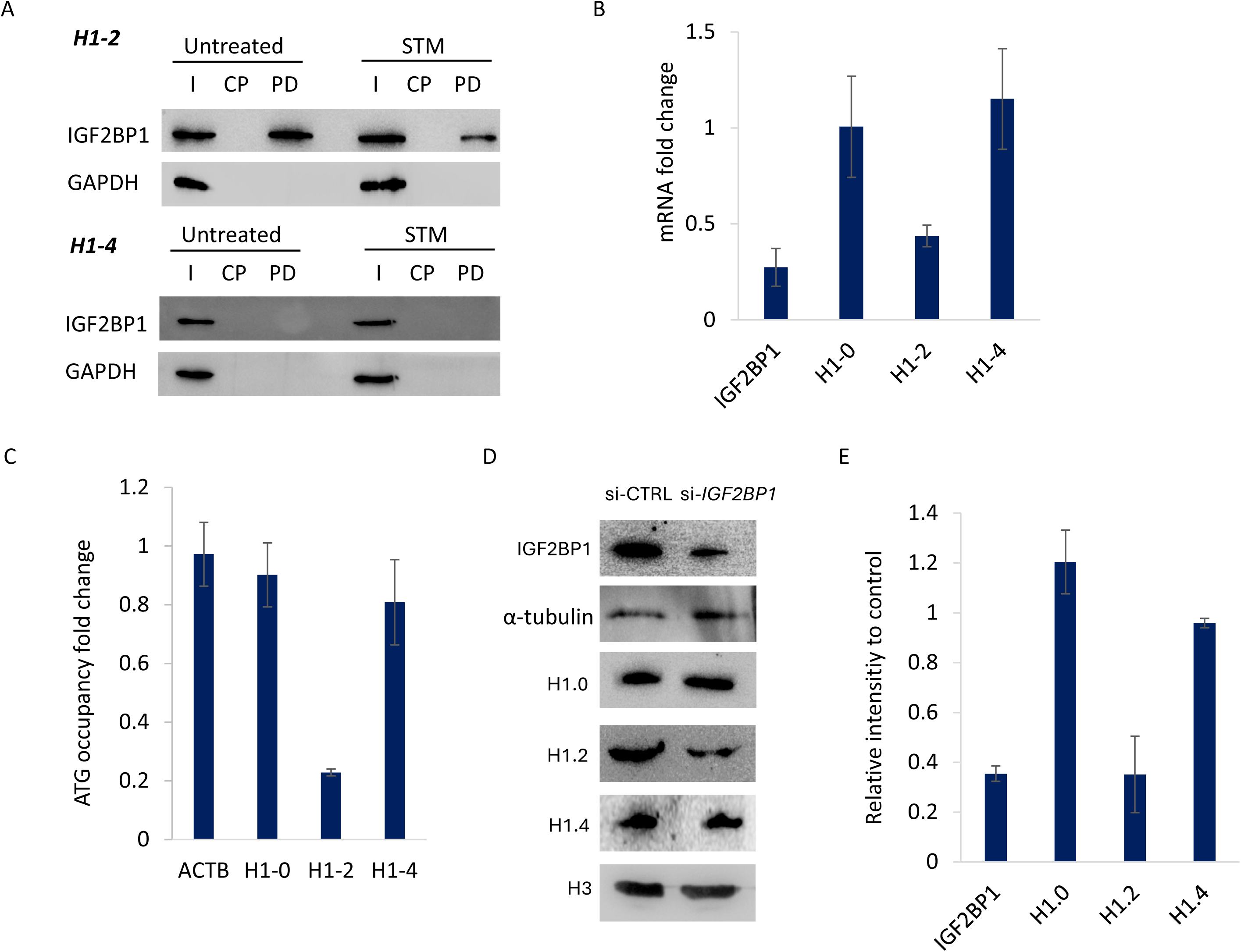
Interaction and function of IGF2BP1 in the regulation of H1 subtypes. A. Western blot against IGF2BP1 after the pull-down (PD) of HeLa cells untreated or treated with 20 µM STM2457 (STM) for 48h, using H1-2 and H1-4 specific probes. I, input. CP, pull-down using a control probe, without specificity for any transcript. PD, pull-down with the specific probe. B, Fold change of the mRNA levels after partial depletion of IGF2BP1 with a specific siRNA. C, Fold change of the transcript levels of the translation start-site region obtained by targeted ribosome profiling. D, Western blots of total protein extracts. E, Quantification of Western blot images. Fold change was calculated respect to the untreated cells (UNTR) or cells transfected with a control siRNA, which were used as a negative control in all the experiments. Error bars correspond to the standard deviation of three biological replicates.

The functional role of IGF2BP1 was confirmed by its partial depletion with a specific siRNA, which reduced the transcript and proteins levels by more than 65% (Figure 5B, 5D, and 5E). IGF2BP1 depletion caused approximately a 50% reduction in the mRNA and protein levels of H1.2, while H1.0 and H1.4 remained unaltered. Ribosome-profiling showed a marked 70% decrease in the ATG occupancy in the *H1-2* mRNA, without changes in the other H1 subtypes (Figure 5C). These results suggest that IGF2BP1 promotes *H1-2* transcript stability and translation in an m6A dependent manner. Three additional m6A readers were identified bound to *H1-2* mRNA, hnRNPA2B1, RBMX (hnRNPG), and PRRC2A, but their functional role in *H1-2* regulation was not studied in this work (13).

The specific proteins found in the pull-down of the *H1-4* transcript were associated with all the enriched GO terms. However, they were especially relevant for mRNA stabilization and the positive regulation of translation, which our experimental data suggest is the functional role of m6A in the regulation of *H1-4* (Figure 4D). We found only two m6A readers bound to *H1-4*, hnRNPD and ELAVL1 (HurR) to a less extent (44–52). The protein hnRNPD was enriched in the *H1-4* pull-down, as observed in the Western blot (Figure 6A). hnRNPD interaction with the *H1-4* transcript seemed strongly dependent on m6A since it was severely reduced in the pull-down performed after m6A inhibition with STM and cycloleucine (Figure 6A and S8B). In the case of *H1-2*, hnRNPD was detected by mass spectrometry, although it was not significantly enriched (Table S6). Surprisingly, the Western blot of the *H1-2* transcript pull-down confirmed this interaction, which also seemed slightly affected after treatment with STM (Figure 6A).

**Figure 6.**
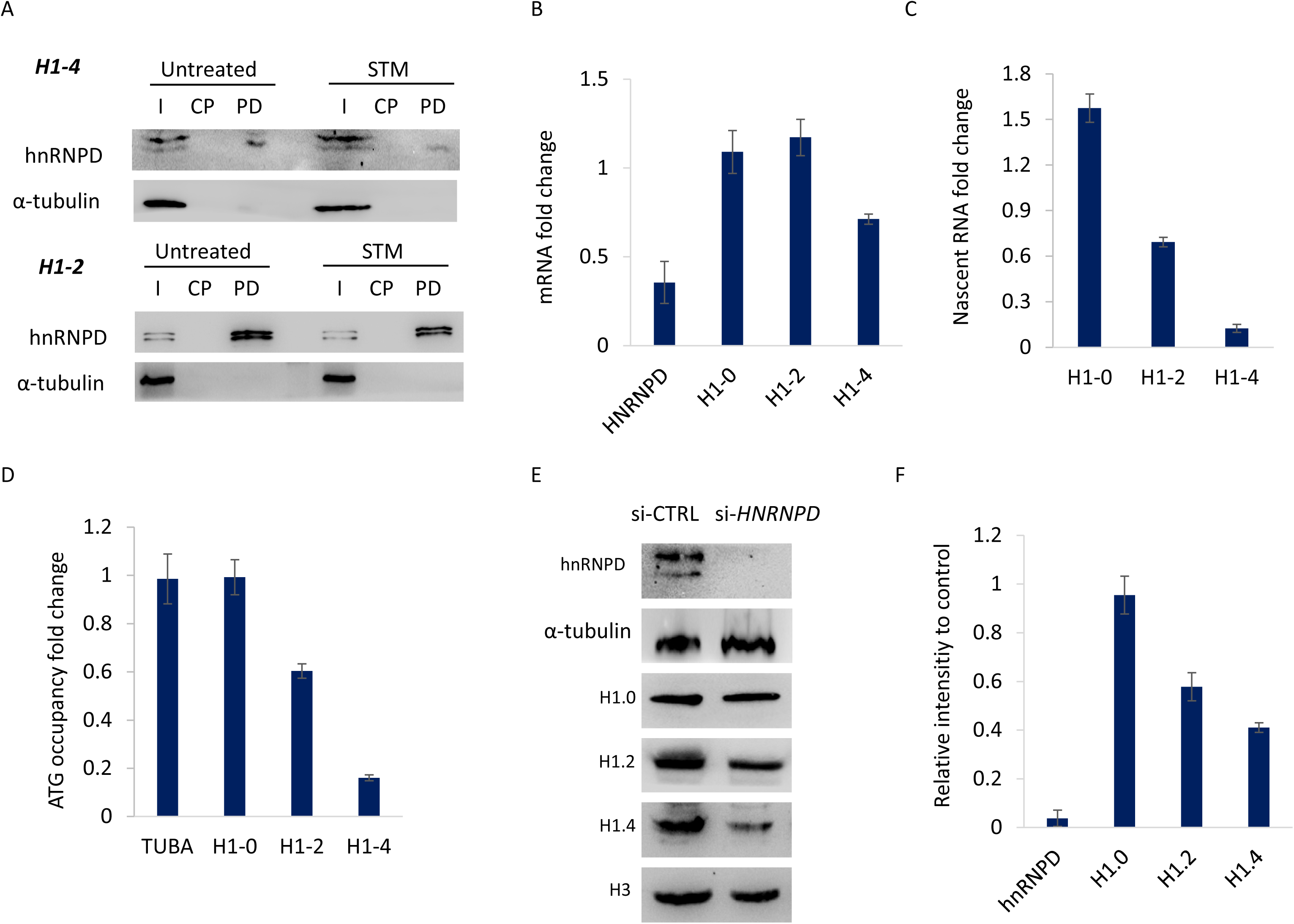
Interaction and function of hnRNPD in the regulation of H1 subtypes. A. Western blot against hnRNPD after the pull-down (PD) of HeLa cells untreated or treated with 20 µM STM2457 (STM) for 48h, using H1-2 and H1-4 specific probes. I, input. CP, pull-down using a control probe, without specificity for any transcript. PD, pull-down with the specific probe. B, Fold change of the mRNA levels after partial depletion of hnRNPD with a specific siRNA. C, Fold change of the transcript levels of the translation start-site region obtained by targeted ribosome profiling. D, Western blots of total protein extracts. E, Quantification of Western blot images. Fold change was calculated respect to the untreated cells (UNTR) or cells transfected with a control siRNA, which were used as a negative control in all the experiments. Error bars correspond to the standard deviation of three biological replicates.

Using a specific siRNA against *HNRNPD* we achieved a depletion of more than 80% of its transcript and the protein was undetectable by Western blot. In these conditions, we observed a 30% decrease in the mRNA levels of *H1-4* and more than 90% reduction in its nascent transcript (Figure 6B and 6C). At the transcript level, no changes were apparent in *H1-0*, while the nascent *H1-2* transcript was reduced to approximately 70%. Ribosome profiling after hnRNPD depletion caused an 80% and 40% reduction in the ATG-occupancy of *H1-4* and *H1-2*, respectively (Figure 6D). As a result, the protein levels of H1.4 showed a 60% decrease and H1.2 40% (Figure 6E and 6F).

In contrast to *H1-2* and *H1-4*, the transcript of *H1-0* has very low levels in HeLa, so the specific pull-down could not identify significantly enriched proteins with FDR < 0.05. Considering that m6A inhibition increased *H1-0* transcript levels without altering de novo transcription, it is reasonable to think that the role of this modification is to promote mRNA degradation. The principal m6A reader associated with mRNA degradation is YTHDF2, so we hypothesized that it might be involved in the regulation of the transcript levels of *H1-0* (13). Additionally, the transcript of *H1-0* has already been identified as bound to YTHDF2 in HeLa by RIP-seq (53).

We performed RNA-immunoprecipitation using an antibody against YTHDF2 and analyzed the presence of several transcripts by RT-qPCR (Figure 7A). The results showed that the *H1-0* transcript was enriched in the immunoprecipitate almost to 15-fold, a value similar to the enrichment of *MYC*, a known target of YTHDF2 (35–37). Treatment with STM significantly decreased the amount of *H1-0* and *MYC* immunoprecipitated, with a reduction of around 30% and 50%, respectively. In contrast, *H1-2*, *H1-4*, and *GAPDH* enrichment in immunoprecipitation was very low in the untreated cells and remained the same in the presence of STM. These results agree with the hypothesis that YTHDF2 binds to m6A in the *H1-0* transcript. YTHDF2 was partially depleted using specific siRNAs, obtaining 80% reduction of its transcript and 70% of the protein (Figure 7B-7D). As expected from the mass spectrometry and RIP results, only *H1-0* was affected with about 3-fold increase in its transcript and protein levels.

**Figure 7.**
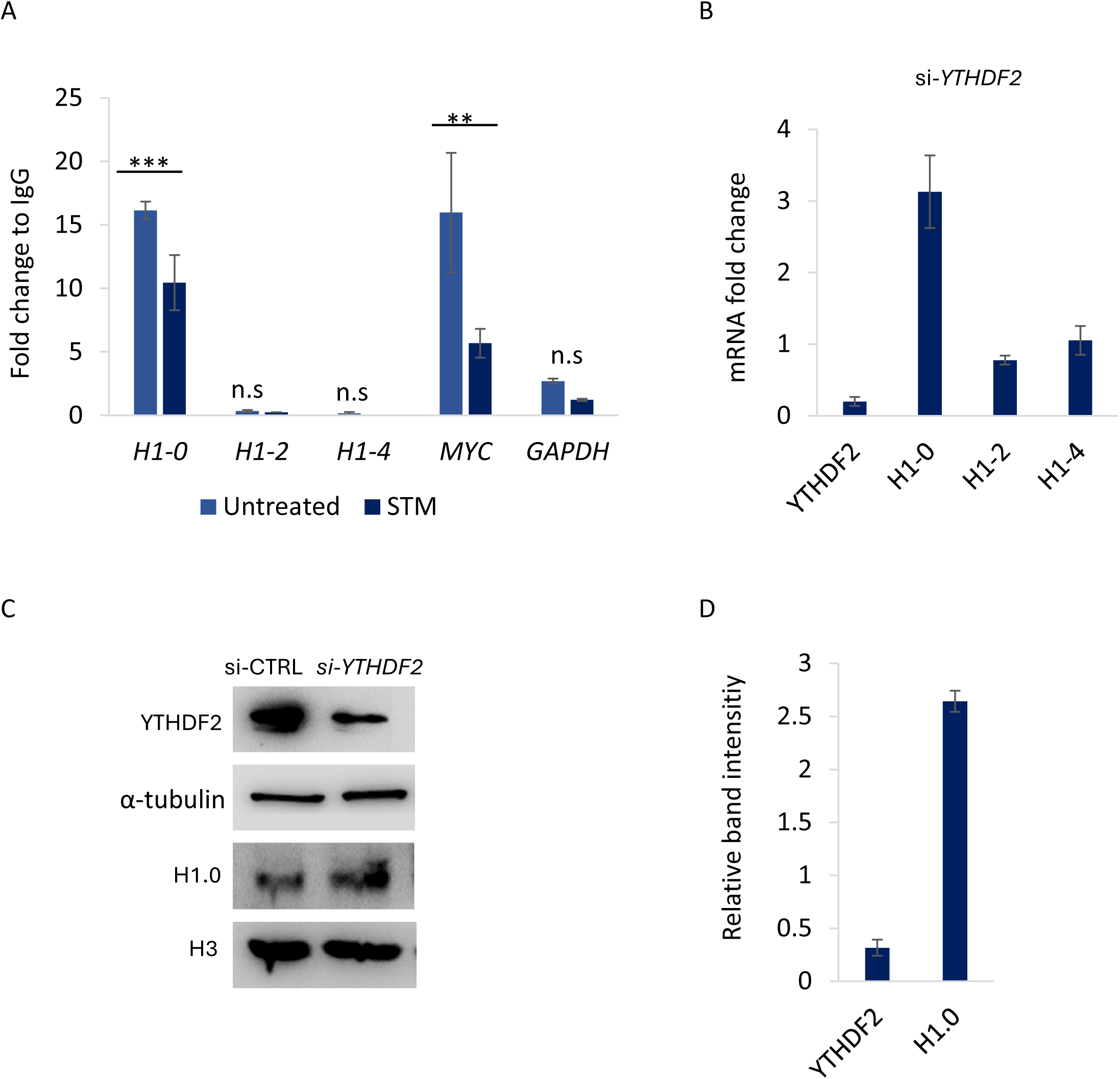
Interaction and function of YTHDF2 in the regulation of H1 subtypes. A. Fold change of the transcript levels after RNA immunoprecipitation with an antibody against YTHDF2 using HeLa cells untreated or treated with 20 µM STM2457 (STM) for 48h. The differences between the mRNA immunoprecipitated in the untreated cells and after treatment with m6A inhibitors were analyzed with a two-tailed Student’s t-test. n.s. not significant. ** p-value < 0.01. *** p-value< 0.001. B, Fold change of the mRNA levels after partial depletion of YTHDF2 with a specific siRNA. C, Western blots of total protein extracts. D, Quantification of Western blot images. Fold change was calculated respect to the untreated cells (UNTR) or cells transfected with a control siRNA, which were used as a negative control in all the experiments. Error bars correspond to the standard deviation of three biological replicates.

## Discussion

The variability of the H1 complement results from the combined action of several regulatory mechanisms. In the last few years, it has been shown that epitranscriptome modifications are involved in the regulation of mRNA metabolism. The most prevalent modification in the mRNA is m6A, which is associated with the processing, transport, stability, and translation of mRNA (12–14). In this work, we studied the contribution of m6A to the regulation of histone H1 subtypes.

We examined the H1 subtypes enriched in m6A in synchronized HeLa cells by a modified meRIP-seq. Considering that the RD H1 subtypes have a short mRNA with approximately 700 bp, we did not fragment the RNA before m6A immunoprecipitation. In this way, we could compare the levels of each transcript before and after the IP and estimate the enrichment in m6A, but we could not localize the modification within the transcript.

Almost 70% of the transcripts with high m6A levels overlapped between cell cycle phases, suggesting that this modification is stable for the most part during the cell cycle. Previous studies in synchronized HeLa cells have reported differences in m6A in G1 and S-phase, but it remained steady from late S to G2/M (53). Both results are not comparable for two reasons. The first reason is that we collected the G1 sample after the first cell division following the release of the double thymidine block, while Fei and colleagues have instead a G1/S sample directly after removing the thymidine block. The second reason is that they compared the m6A peaks using fragmented RNA. Instead, we analyzed if each transcript was enriched in the m6A immunoprecipitation, comparing its levels with those in the RNAseq in the same phase.

Regarding histone H1 subtypes, we found that the five subtypes expressed in HeLa had different levels of m6A. Subtypes *H1-0* and *H1-4* have high m6A levels, and *H1-2* has medium levels of this modification. Our experimental approach did not allow the localization of m6A within the transcripts. However, the information in the RMBase 3.0 suggests that m6A is probably located in the coding region of *H1-2* and *H1-4* and in the 3’UTR of *H1-0* (22). The latter was confirmed by eTAM-seq in HeLa cells, which provides single-nucleotide resolution to m6A localization (38). Interestingly, despite the H1 subtypes being differentially expressed during the cell cycle, the levels of m6A appeared to be constant, except for *H1-0* in G2/M (9). These results suggest that the effects of m6A in *H1-2* and H1-4 are not directly linked to the fluctuations during the cell cycle, whereas in *H1-0*, it seems to contribute to the changes from G2/M to G1.

To study the role of m6A in the regulation of H1.0, H1.2, and H1.4, we analyzed the impact of m6A inhibition in several processes and the readers involved in its recognition. To address the effect of m6A, we examined the mRNA and protein levels, de novo transcription, and ribosome occupancy. To identify m6A readers, we used specific mRNA pull-down coupled with proteomics and RIP coupled to RT-qPCR.

We observed that the decrease in m6A caused an increase in the mRNA and protein levels of H1.0 using two different m6A inhibitors. No alterations in transcription or translation were detected. These results suggested that the role of m6A is associated with mRNA degradation (Figure 8). The mRNA levels of *H1-0* in HeLa are low, which prevented the use of a proteomic-based approach for reader identification. Considering that m6A promotes *H1-0* mRNA degradation, that most m6A sites are in the 3’UTR, and previous knowledge of m6A readers, we hypothesized that YTHDF2 could be binding m6A in the *H1-0* transcript (13). Using RIP coupled with RT-qPCR, we found that the mRNA of *H1-0* was bound to YTHDF2 and that this interaction significantly decreased upon m6A inhibition with cycloleucine, as well as STM2457, which confirmed observations from a previous study that identified the *H1-0* transcript as a target of YTHDF2.

**Figure 8.**
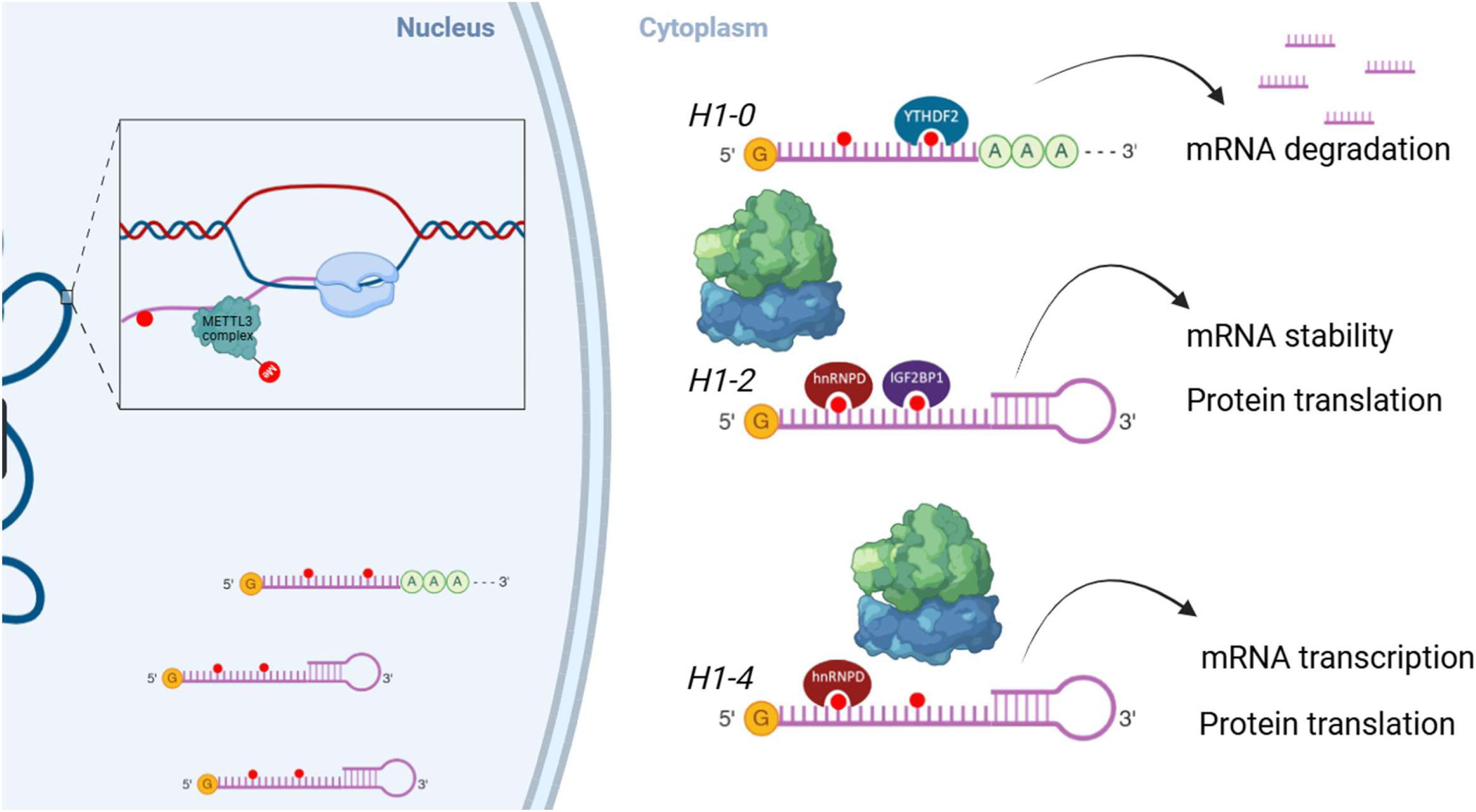
Schematic model of the role of m6A in the regulation of H1 subtypes.

Furthermore, the YTHDF2 protein accumulates in G2/M, which could explain the decrease in the m6A levels of *H1-0* in this phase of the cell cycle and the downregulation of its transcript levels (53). The functions of the different members of the YTHDF family seem to be redundant in HeLa cells (54). However, considering that the partial depletion of YTHDF2 reproduced the results obtained by m6A inhibition, it seems unlikely that other family members play a significant role in its regulation.

Inhibition of m6A triggered a decrease in the mRNA and protein levels of *H1-2* and in the occupancy of the translation start by the ribosome, without major alterations of the levels of the nascent transcript. These data suggest that m6A promotes the stability and translation of *H1-2* mRNA (Figure 8). This hypothesis agrees with the identification by proteomics of IGF2BP1 and IGFBP2 specifically bound to the *H1-2* transcript, which is associated with mRNA stability and negative regulation of translation (12–14). We detected IGF2BP1 interaction with *H1-2* mRNA by RNA-pull-down followed by Western blot, which decreased upon m6A inhibition, confirming that the interaction was mediated by this modification. The functional role of this m6A reader in the regulation of *H1-2* mRNA stability and translation in HeLa cells was validated by its partial depletion. We also detected the interaction of *H1-2* transcript with hnRNPD by RNA-pull-down followed by Western blot, although in this case the interaction showed very little changes upon m6A inhibition. Partial depletion of hnRNPD reduced *H1-2* ATG-occupancy by the ribosome, suggesting a role in its translation.

Three more m6A readers were identified in the *H1-2* transcript pull-down. Two belong to the heteronuclear ribonucleoproteins, hnRNPA2B1 and RBMX (hnRNPG). hnRNPA2B1 promotes m6A-dependent primary-microRNA processing and alternative splicing (55). It can also be involved in the 3’-end processing and the regulation of mRNA stability and translation, independent of m6A (56). hnRNPG m6A-dependent functions are associated with mRNA stability and alternative splicing (57). Different roles have been assigned to PRRC2A in specific cell types. In neural cells, it stabilized critical transcripts, promoting oligodendrocyte progenitor cell proliferation and oligodendrocyte fate determination (58). In mice testes, PRRC2A reduced the abundance and promoted translation of its targets, regulating the transition of the expression profile from spermatogonia to spermatocytes and the progression of male meiotic metaphase (59). Although functions associated with RNA splicing are discarded, as H1 genes are intron-less, future studies will determine whether these readers contribute to the processing and stability of the *H1-2* transcript.

Proteomic results showed that the m6A reader hnRNPD was significantly enriched in the *H1-4* mRNA pull-down, and its presence was confirmed by Western blot. Interaction of hnRNPD with *H1-4* transcript seems to depend on m6A as the protein was strongly reduced in the pull-down after treatment with STM and cycloleucine. Several functions have been described for hnRNPD: i) It decreased the stability of several m6A-modified transcripts, *YAP1, ATF3*, and *TFEB* (44–46); ii) It promoted *ZEB1-AS1* and *MYC* translation without altering the mRNA levels(47, 48, 52); iii) It reduced the stability and promoted the translation of *CPEB1* (49). In the *H1-4* transcript, we observed that m6A inhibition resulted in almost no changes in its mRNA levels. In contrast, the nascent transcript and the ribosome occupancy of the translation start site were severely reduced, which decreased the protein levels (Figure 8). The partial knockdown of hnRNPD confirmed its role in the regulation of H1.4 protein levels.

The lack of effect of hnRNPD on *H1-4* transcript stability could be explained in two ways. First, by ELAVL1, the other m6A reader found in the *H1-4* pull-down, which promotes *SOX2* and *MALAT1* mRNA stability in gliomas (50, 51). Second, by the reduction in *H1-4* de novo transcription. Considering the reduction in the ribosome occupancy, we expected a higher decrease in the protein levels. Perhaps an increase in protein stability might explain the final H1.4 protein levels.

We found several proteins associated with stress granule assembly in the pull-downs of *H1-2* and *H1-4* transcripts, including TIA1, CSDE1, G3BP1, and G3BP2 (60). Discrepant relationships between m6A and stress granules have been described. On the one hand, it has been reported that m6A inhibits G3BP1 and G3BP2 binding to mRNA (61). On the other hand, YTHDF2 bound to mRNA containing multiple m6A undergo liquid-liquid phase separation and participates in the formation of stress granules, while IGF2BP1 stabilizes mRNA within them (62–64). Additionally, hnRNPD is also recruited to stress granules upon several stimuli (65). It seems that *H1-2* and *H1-4* transcripts are targeted to stress granules either by m6A modification or by direct binding to SG-core proteins. Future studies would be needed to understand the role of stress granules in the regulation of H1 transcripts, as well as that of other epitranscriptome modifications such as m5C.

In summary, we described, for the first time, the differential regulation of several H1 subtypes by m6A (Figure 6). It promotes the degradation of the *H1-0* transcript through the interaction with YTHDF2, presumably at the 3’UTR. In the case of *H1-2*, m6A is recognized by IGF2BP1, enhancing the stability and translation of its transcript. For *H1-4*, m6A facilitates the binding of hnRNPD, which favors the transcription and translation of this subtype. Our findings revealed a new piece of the complex regulation of H1 subtypes, which may contribute to the variability of the H1 complement in health and disease.

## Data Availability

The data underlying this article are available in the article and in its online supplementary material. RNA-seq data (Supplementary Tables S3 and S4) were deposited in the Gene Expression Omnibus database under the accession number GSE236863. Proteomic data (Supplementary Tables S6 and S7) were deposited at the PRIDE repository under the identifier PXD043530.

## Author Contributions Statement

Conceptualization: Ponte I, Jordan A, and Roque A. Formal analysis: García-Gomis D, López-Gómez J, Serna-Pujol N. Funding acquisition: Jordan A and Roque A. Investigation: García-Gomis D, López-Gómez J, Andrés A, Fernández Z, and Benyahya A. Supervision: Ponte I, Jordan A, and Roque A. Visualization: García-Gomis D, López-Gómez J, and Roque A. Writing – original draft: García-Gomis D, López-Gómez J, and Roque A. Writing – review & editing: Ponte I, Jordan A, and Roque A

