## Supplementary figures for "Differential regulation of histone H1 subtypes by N6-methyladenosine RNA methylation"

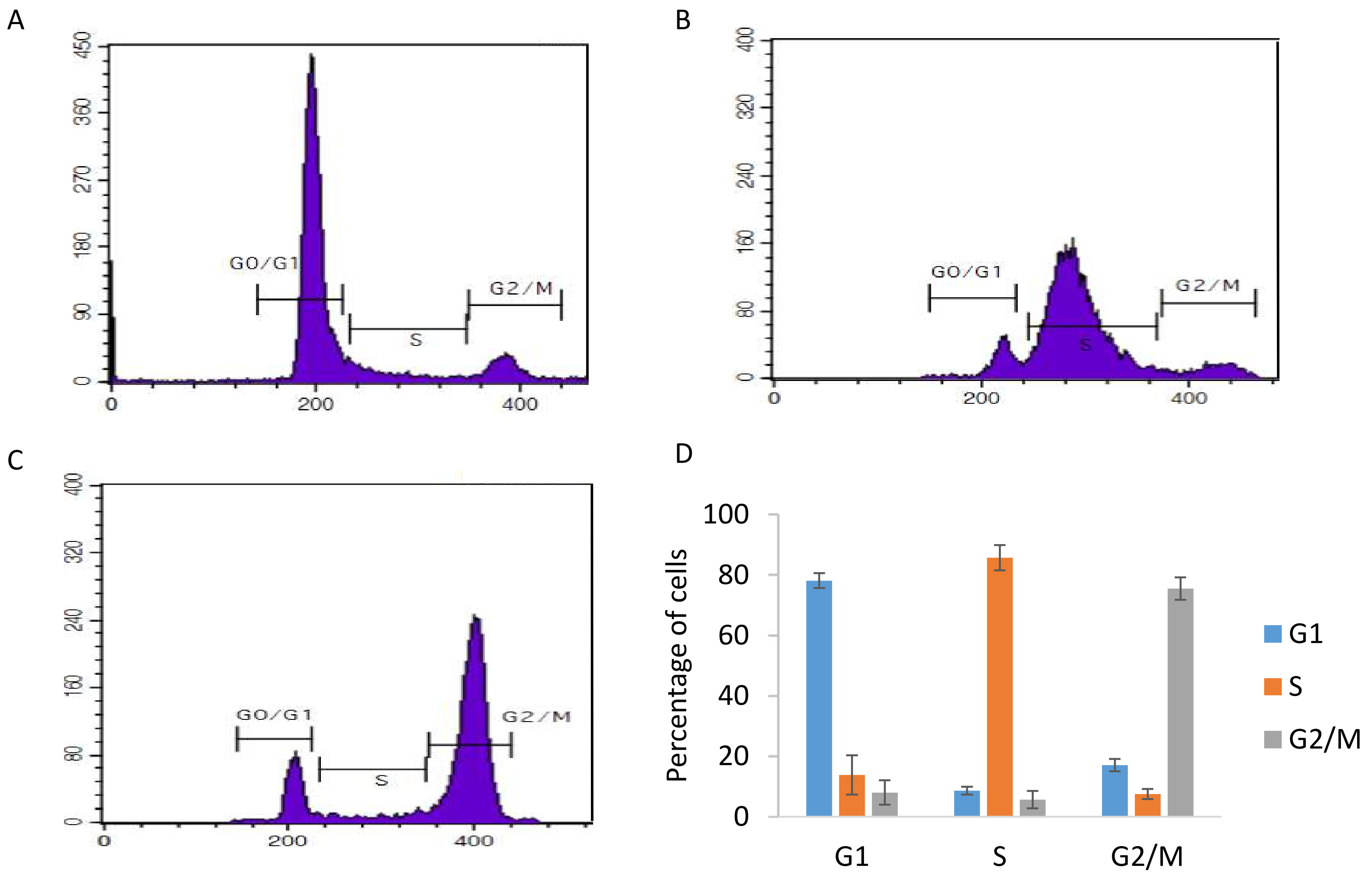


**Figure S1. Synchronization of HeLa cells**. Representative flow cytometry profiles of HeLa cells synchronized in G1 (panel A), S (panel B), and G2/M (panel C). D. Quantification of the synchronized samples. Error bars correspond to the standard deviation of three biological replicates.

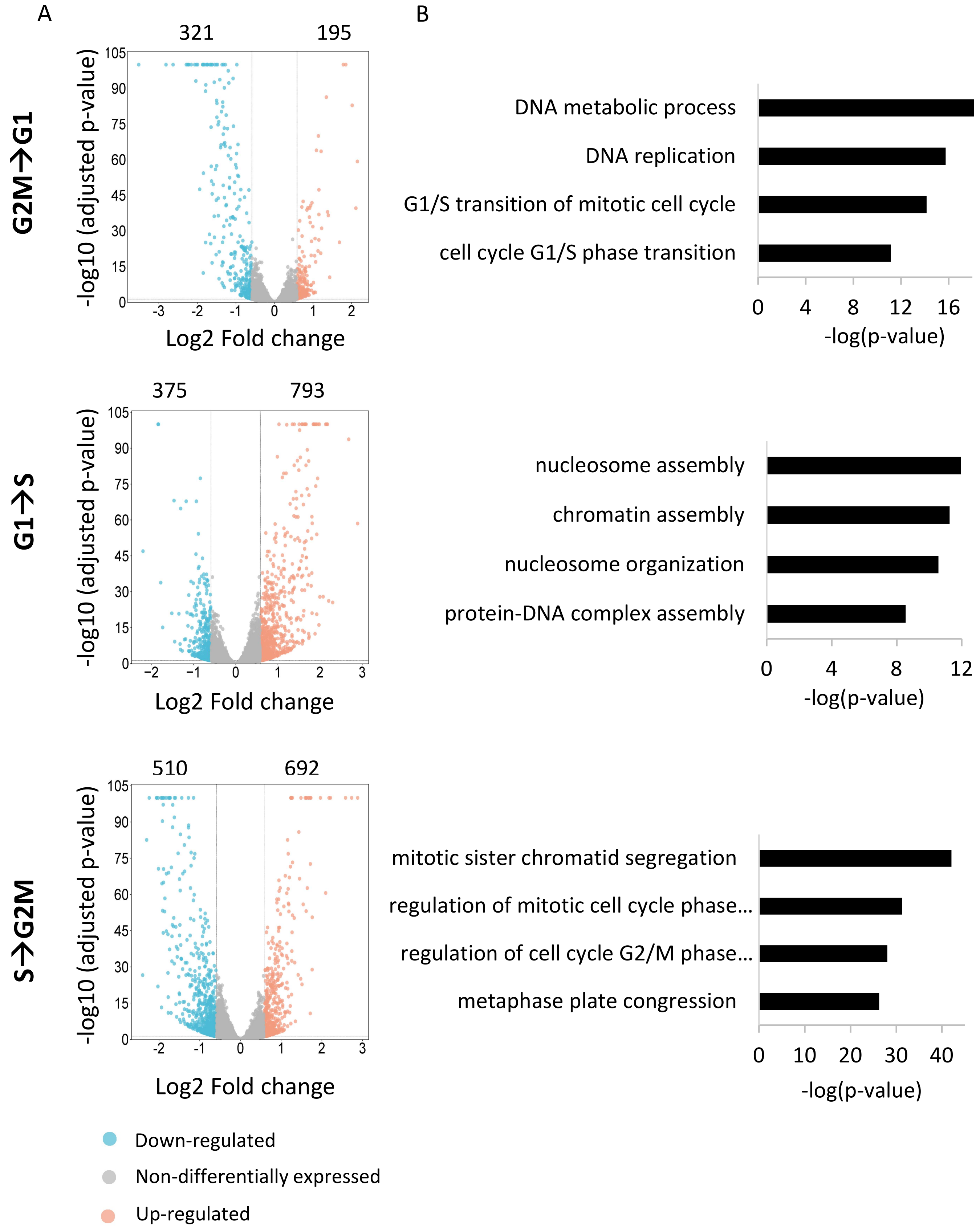


**Figure S2. Gene expression during cell cycle.** A. DESeq2 differential gene expression analysis of synchronized HeLa cells. Genes were considered upregulated when their transcripts had a fold change >1.5 and FDR-adjusted p-value < 0.05. Genes were considered downregulated when their transcripts had a fold change <1/1.5 and FDR-adjusted p-value < 0.05. The number of upregulated or downregulated genes in each cell cycle transition is indicated. B. GO term analysis showing the biological processes enriched in the upregulated genes in each phase of the cell cycle.

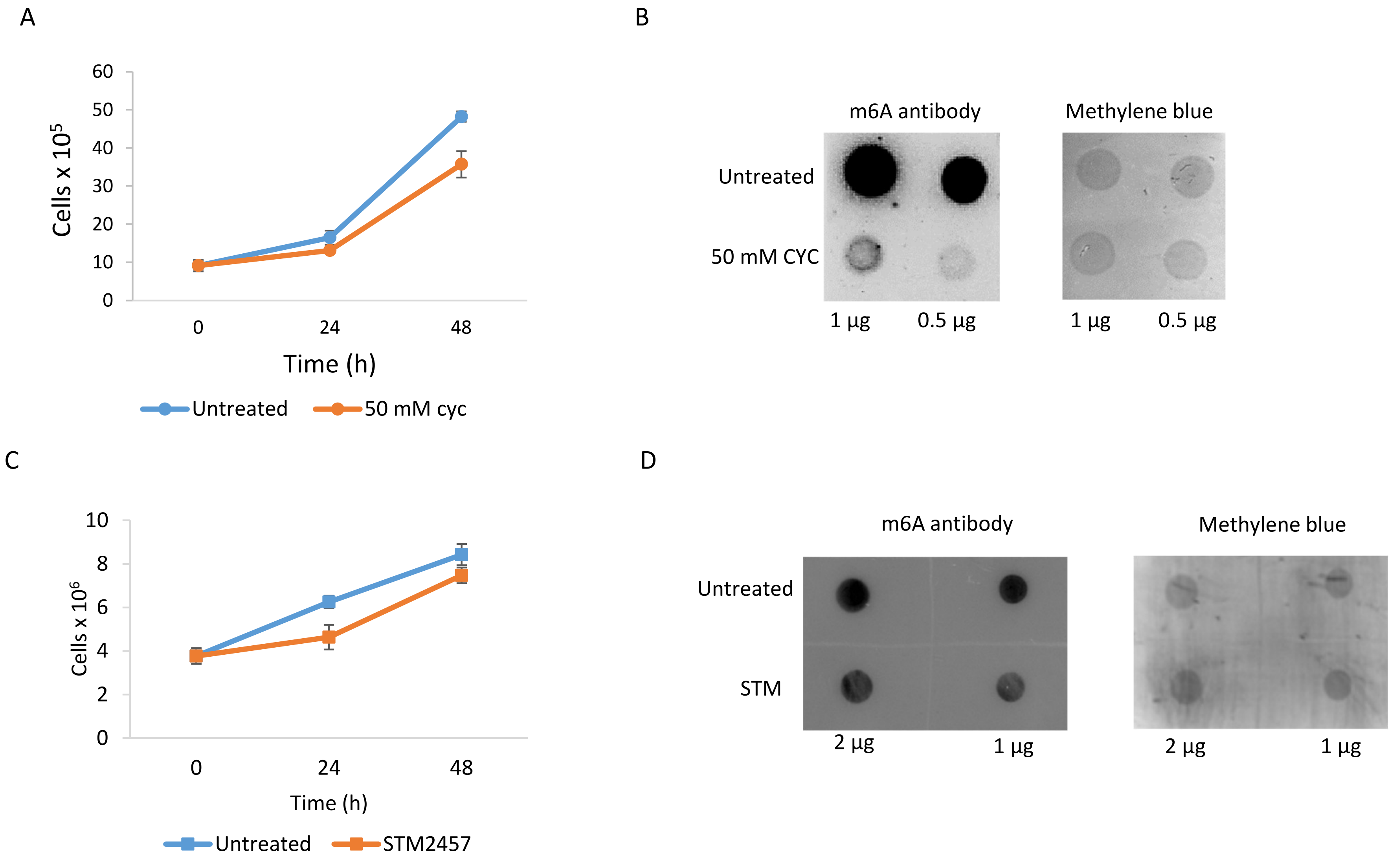


**Figure S3. Effects of m6A inhibition in HeLa.** A, C. Growth curve of HeLa cells treated with 50 mM cycloleucine (CYC) for 24h or with 20 µM STM2457 (STM) for 48h. B, D. Dot blot against m6A (left) and loading control stained with methylene blue (right). Error bars correspond to the standard deviation of three biological replicates. Untreated cells were used as a negative control.

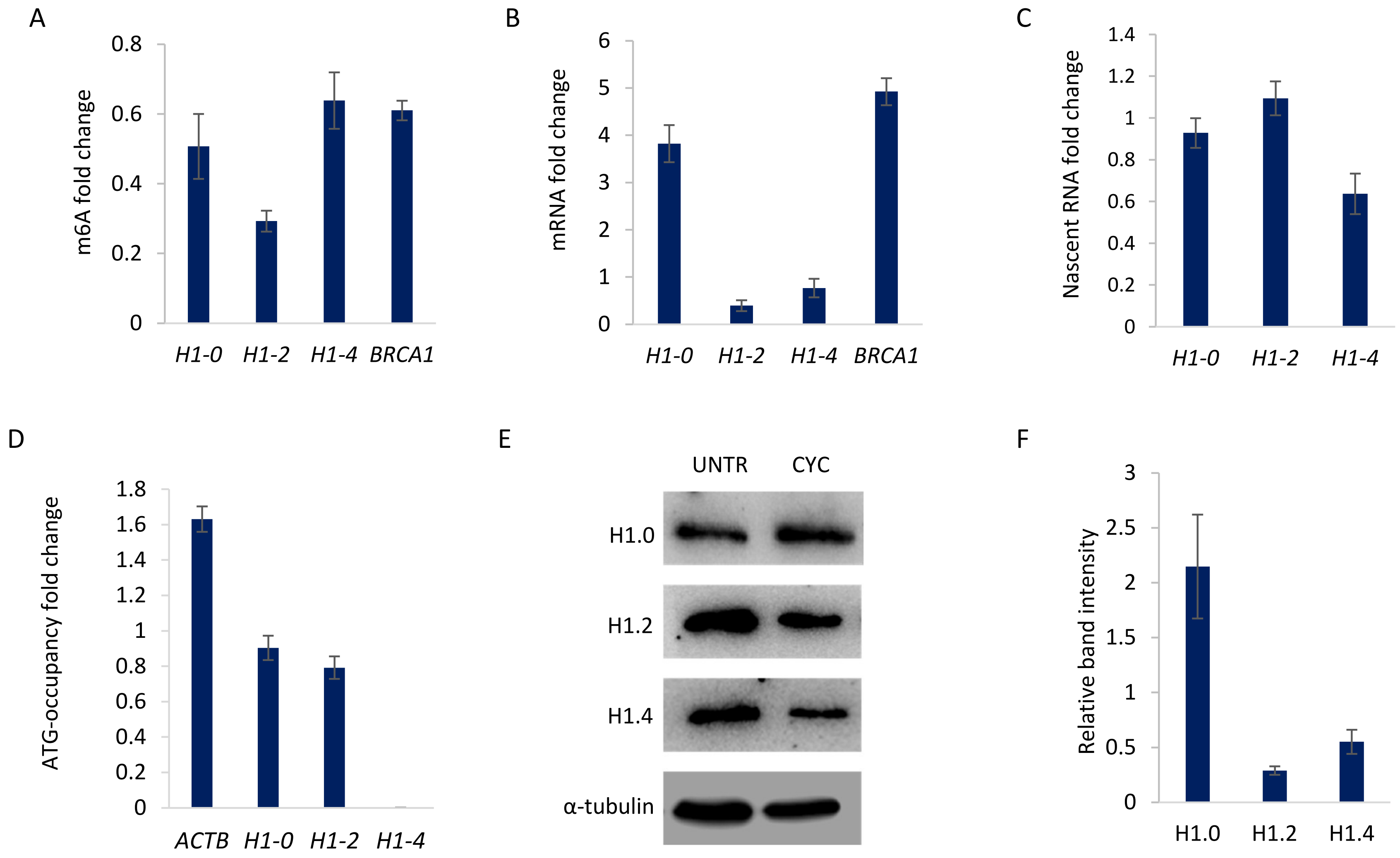


**Figure S4. Effects of m6A inhibition by treatment with cycloleucine (CYC).** HeLa cells were treated with 50 mM CYC for 24h. Fold change was calculated respect to the untreated cells (UNTR), which were used as a negative control in all the experiments. A. Fold change of the m6A-immunoprecipitated mRNA. B. Fold change of the mRNA levels. C. Fold change of the BrdU-labeled nascent RNA immunoprecipitated after nuclear run on. D, Fold change of the transcript levels of the translation start site region obtained by targeted ribosome profiling. E, Western blots of total protein extracts. F, Quantification of the Western blot images. Error bars correspond to the standard deviation of three biological replicates.

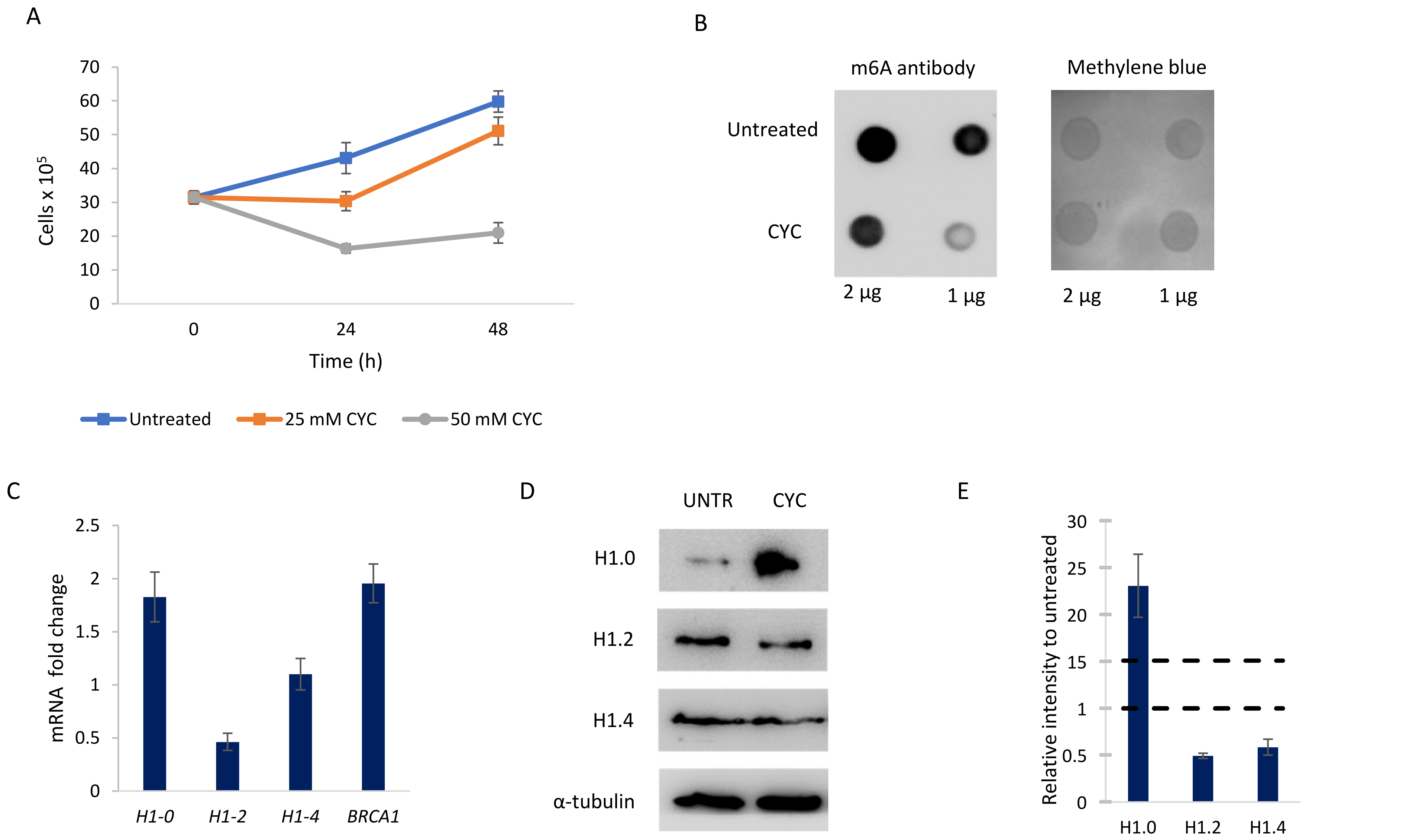


**Figure S5. Effects of m6A inhibition in HEK293T cells.** A. Growth curve of HEK293T cells treated with cycloleucine (CYC). B. Dot blot against m6A (left) and loading control stained with methylene blue (right). C. mRNA fold change of the cells treated with both inhibitors, with respect to the control. D. Western blots of total protein extracts after m6A inhibition. E. Quantification of the Western blot images, relative to control and normalized by tubulin. Unless specified in the figure, HEK293T cells were treated with 25 mM cycloleucine for 24h. Error bars correspond to the standard deviation of three biological replicates. Untreated cells were used as a negative control. Horizontal black dashes represent truncation in the y-axis.

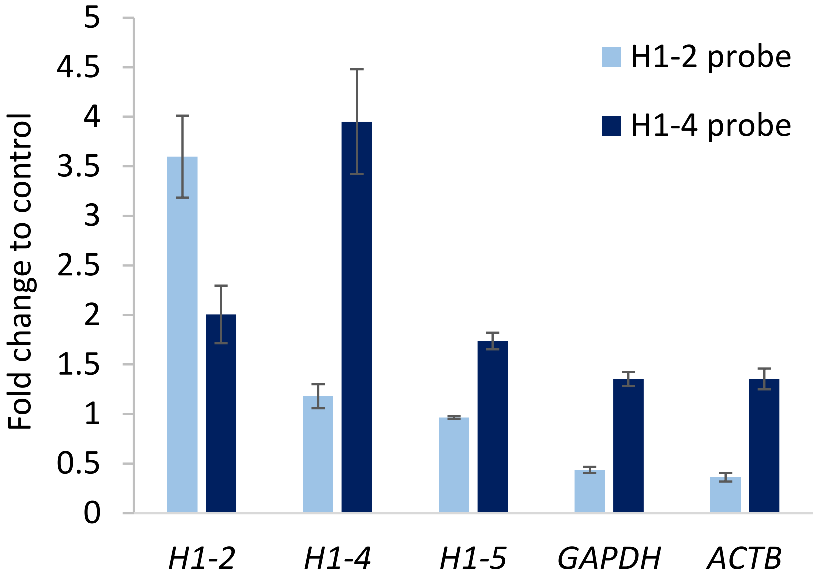


**Figure S6. Analysis of probe specificity for mRNA pull-down.** Fold change to the control probe of the pull-down using *H1-2* and *H1-4* specific probes obtained by RT-qPCR. Error bars correspond to the standard deviation of the triplicates.

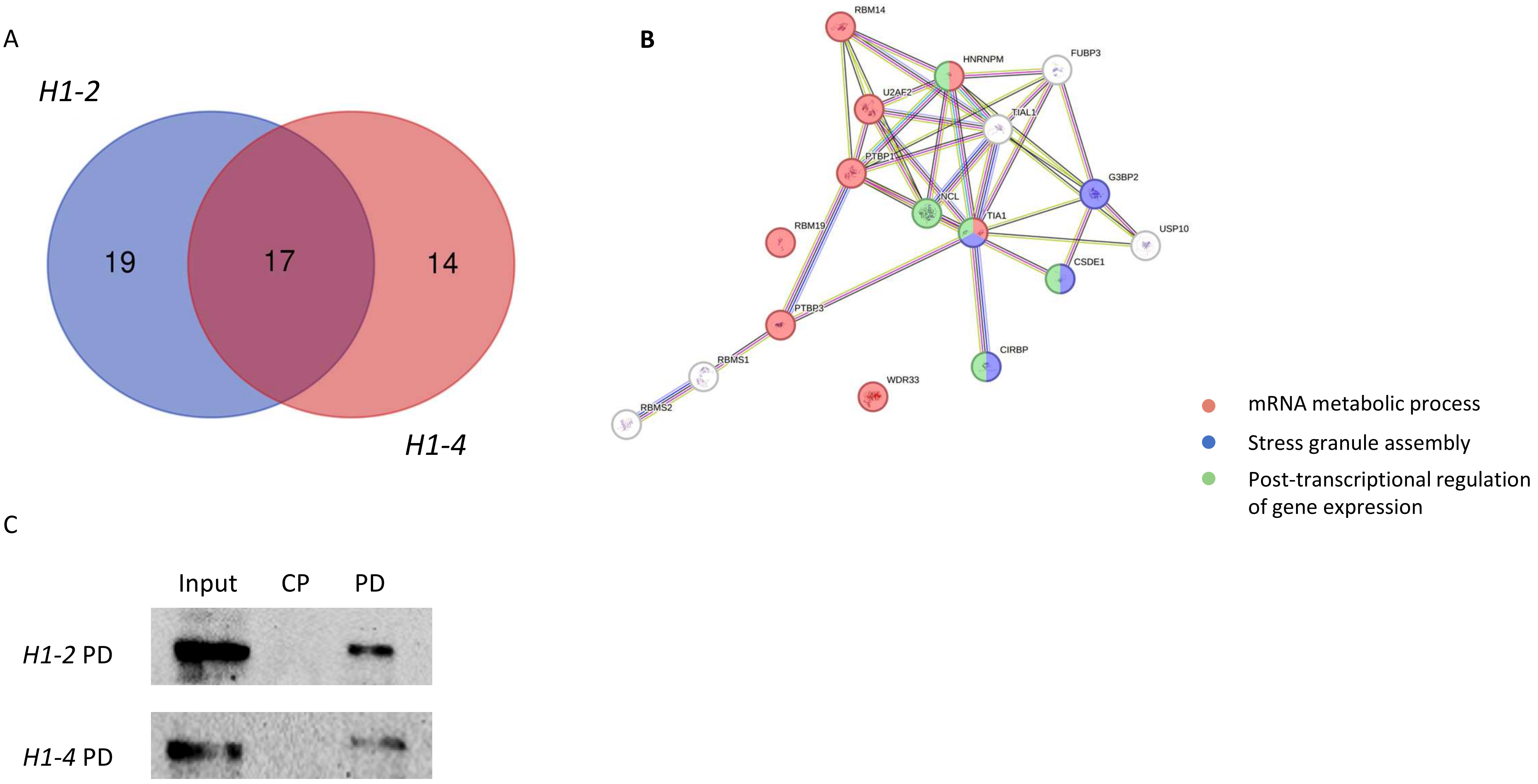


**Figure S7. Common proteins bound to *H1-2* and *H1-4* transcripts.** A. Venn diagram of the proteins identified in the pull-downs of *H1-2* and *H1-4* transcripts by mass spectrometry. B. String interaction network of the proteins common in *H1-2* and *H1-4* pull-downs. The color in the nodes denotes their role in the biological processes shown in the legend. C. Western blot against hnRNPM of the *H1-2* and *H1-4* pull-down (PD). CP, control probe.

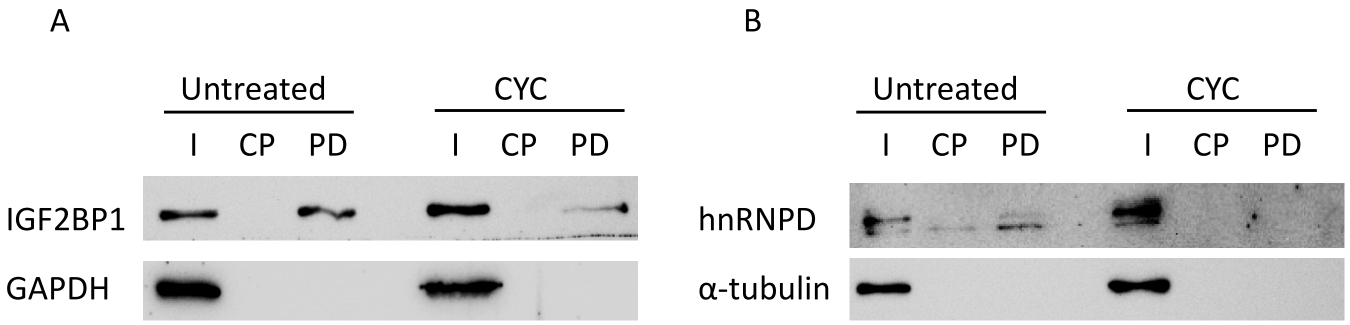


**Figure S8. Effect of cycloleucine on the binding of the main m6A readers to *H1-2* and *H1-4* transcripts. A,** Western blot against IGF2BP1 after the pull-down (PD) of HeLa cells untreated or treated with cycloleucine (CYC) using *H1-2* specific probe. B, Western blot against hnRNPD after the pulldown of HeLa cells untreated or treated with cycloleucine (CYC) using *H1-4* specific probe. HeLa cells were treated with 50 mM cycloleucine for 24h. I, input. CP, pull-down using a control probe, without specificity for any transcript. PD, pull-down with the specific probe.
